# PD-1 Mediated Regulation of Unique Activated CD8^+^ T Cells by NK Cells in the Submandibular Gland

**DOI:** 10.1101/2023.09.15.557930

**Authors:** Samantha M. Borys, Shanelle P. Reilly, Ian Magill, David Zemmour, Laurent Brossay

## Abstract

The increasing utilization of anti-PD-1 immune checkpoint blockade (ICB) has led to the emergence of immune-related adverse events (irAEs), including sicca syndrome. Interestingly, we found that the submandibular gland (SMG) of PD-1 deficient mice harbors a large population of CD8^+^ T cells, reminiscing ICB induced sicca. This phenotype was also observed in the SMG of both NK cell-depleted C57BL/6 animals and NK cell-deficient animals. Mechanistically, using mice conditionally deficient for PD-L1 in the NK cell lineage, we discovered that NK cells regulate CD8^+^ T cell homeostasis via the PD-1/PD-L1 axis in this organ. Importantly, single-cell RNA sequencing of PD-1 deficient SMG CD8^+^ T cells reveals a unique transcriptional profile consistent with TCR activation. These cells have limited TCR diversity and phenotypically overlap with GzmK^+^ CD8^+^ T autoimmune cells identified in primary Sjögren’s syndrome patients. These insights into NK cell immunoregulation in the SMG, and the consequences of disrupted CD8^+^ T cell homeostasis, provide opportunities for preventing the development of irAEs.

**Highlights:**

- Elevated CD8^+^ T cells in the submandibular gland (SMG) of PD-1 deficient mice parallel sicca-like irAEs seen in ICB patients.
- In addition to their previously described hyporesponsive phenotype, NK cells in the SMG regulate CD8^+^ T cell homeostasis through the PD-L1/PD-1 axis.
- PD-1 deficient SMG CD8^+^ T cells display unique transcriptional profiles associated with proinflammatory functions, TCR activation, interferon stimulation, and exhaustion.
- Oligoclonal expansion and similarities in TCR sequences indicate T cell activation and a preference for recognizing specific antigens.

## Introduction

The submandibular gland (SMG) is host to several tissue-specific lymphocyte subsets, including natural killer (NK) cells and T cells. Residing within non-lymphoid tissues, specialized subsets of tissue-resident NK cells (trNK) help to protect their unique respective tissues. NK cells of the SMG exhibit features of tissue residency and are phenotypically unique^1-4^. Additionally, the SMG is home to a modest population of CD8^+^ T cells, many of which display a tissue-resident memory phenotype^5,6^. Several bacteria and viruses, particularly β- and γ-herpesviruses, infect the SMG, highlighting the importance of tissue-resident immunosurveillance in this organ^7-9^. Interestingly, previous work from our lab and others describes the hyporesponsiveness of NK cells^10^ and T cells^11^ in the SMG. This hyporesponsive microenvironment suggests that there are mechanisms within the SMG to help maintain localized tolerance. Patient data suggests that tolerance in the SMG is broken during immune checkpoint blockade (ICB). One of the most well-studied immune checkpoints, programmed cell death 1 (PD-1), has become a prominent therapy in oncology over the past decade. Immune checkpoints, including PD-1, balance the immune response during infections to prevent immune-mediated damage and protect tissue function^12,13^. Therapeutic blockade of PD-1, or its ligand PD-L1, can be harnessed to restore immune activation to eliminate tumor cells, and has been used to treat a variety of cancers including head and neck cancer, urothelial cancer, colorectal cancer, non-small-cell lung cancer, melanoma, and renal-cell carcinoma^14^. While extremely beneficial for killing tumor cells, PD-1 blockade has been shown to inadvertently cause activation of aberrant T cells, inducing autoimmunity in some patients^15^. As PD-1 blockade becomes a more frequently relied upon cancer therapy, so is the prevalence of these immune-related adverse events (irAEs). Patients treated with PD-1 blockade can develop various irAEs including sicca syndrome^16^, vitiligo^17^, and nephritis^18^. Additionally, loss of PD-1 in mice has been reported to induce autoimmunity including nephritis^13^, cardiomyopathy^19^, and diabetes^20^. Furthermore, PD-1 deficiency in some autoimmune-prone mouse backgrounds can lead to the development of sicca syndrome^13,21^. Sicca syndrome is a chronic systemic autoimmune disease characterized by dryness of eyes and mouth which can cause tooth decay, as well as difficulty speaking, swallowing, and wearing dentures^22^.

Similar to phenotypes observed following PD-1 blockade therapies in humans, we found that the SMG of PD-1 deficient animals are home to a large number of CD8^+^ T cells not detected in wildtype animals. Using a variety of approaches, we discovered that the hyporesponsive salivary gland NK cells act as regulatory cells in this organ. In the absence of NK cells and/or when deficient for PD-L1, CD8^+^ T cells expand dramatically in this organ. Using single-cell RNA sequencing and single cell TCR sequencing techniques, we found that these CD8^+^ T cells are marked with a highly activated transcriptional profile, have limited TCR diversity, and phenotypically overlap with GzmK^+^ CD8^+^ T autoimmune cells identified in humans with primary Sjögren’s syndrome.

## Results

### Expansion of CD8^+^ T cells in SMG of PD-1^-/-^ mice

We and other groups previously reported that NK cells^10^ and CD8^+^ T cells^11^ of the murine SMG are hyporesponsive. Importantly this hyporesponsive phenotype is not intrinsic, and instead depends on the salivary gland environment^2^. We therefore hypothesized that inhibitory receptors regulate these subsets of cells *in situ*. While testing a variety of inhibitory receptor-deficient animals, we noticed that salivary glands from naive PD-1 deficient animals harbor a large proportion of CD8^+^ T cells, not detected in littermate animals **(Figure 1A)**. The PD-1 deficient animals were generated using CRISPR/Cas9 mediated deletion of exon 2 of *Pdcd1*, directly on C57BL/6 (B6) zygotes (**Supplemental Figure 1A**). The resultant line has a 632 bp deletion in exon 2 and flanking introns and completely lacks PD-1 expression at the cell surface, compared to PD-1^+/+^ and PD-1^+/-^ littermate controls (**Supplemental Figure 1B**). In wildtype (WT) C57BL/6 mice, lymphocytes of the SMG are normally composed of a majority of NK cells, with a small population of CD4^+^ and CD8^+^ T cells, and very little to no B cells **(Figure 1A).** As mentioned above, SMG of PD-1^-/-^ mice exhibit a marked increase in frequency **(Figure 1B and Supplemental Figure 1D)** and number **(Figure 1C)** of CD8^+^ T cells. In contrast, CD4^+^ T cell frequency and number are not significantly enriched compared to littermate wildtype controls **(Figure 1D and 1E)**. This phenotype was recapitulated in PD-1^-/-^ mice obtained from vendors, ruling out a potential CRISPR/Cas9 off-target effect **(Supplemental Figure 1C)**. Importantly, although the overall number of CD8^+^ T cells in both wildtype and PD-1^-/-^ SMG is higher in males due to natural sexual dimorphism of the murine salivary gland, the frequency of CD8^+^ T cells increases similarly in both males and females, indicating a lack of sexual dimorphism in the PD-1 deficient phenotype **(Supplemental Figure 1E and 1F)**.

**Figure 1:**
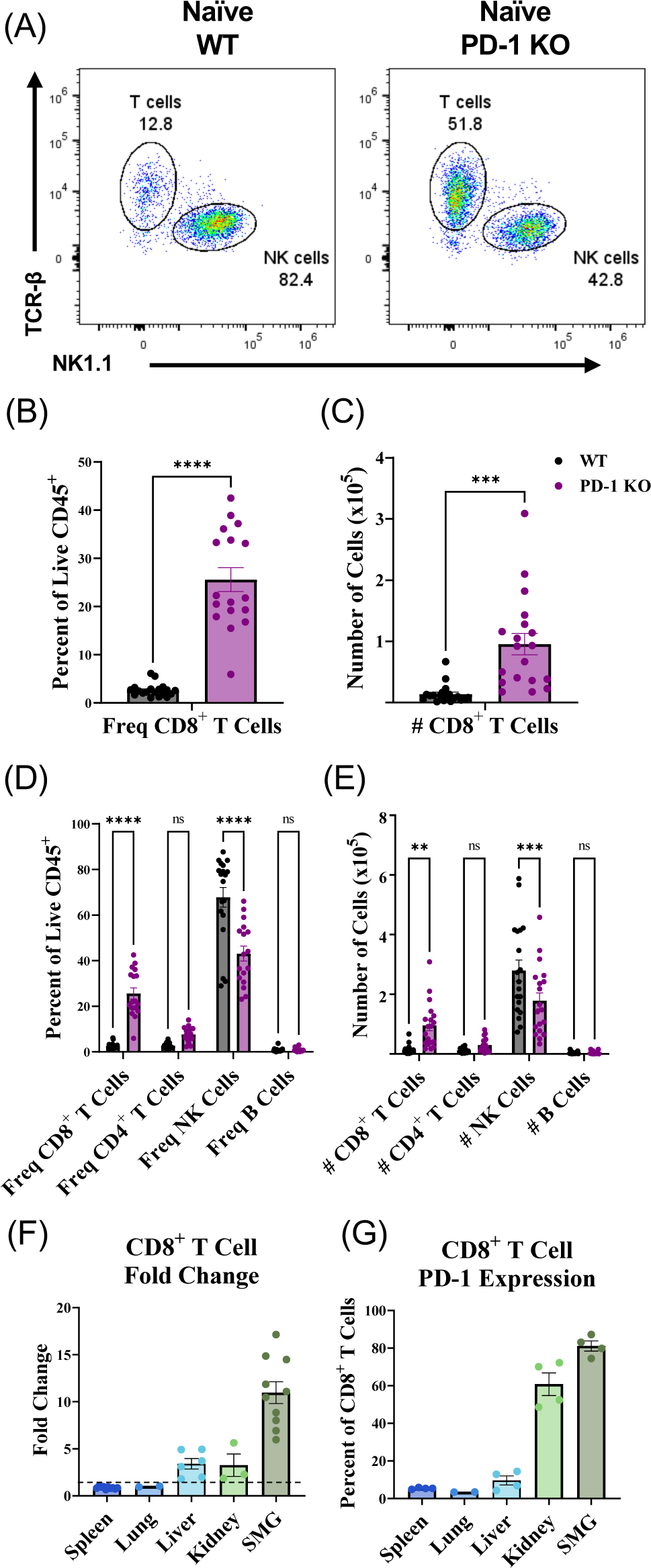
SMG of PD-1 KO mice display elevated frequency and number of CD8^+^ T cells. (A) Representative flow cytometry staining showing distribution of T cells and NK cells in WT vs PD-1 KO SMG, gated on live CD45^+^ lymphocytes. (B) CD8^+^ T cell frequency and (C) number in WT vs PD-1 KO SMG. (D) CD8^+^ T cell, CD4^+^ T cell, NK cell, and B cell frequency and (E) number in WT vs PD-1 KO SMG. Data pooled from 6 independent experiments, n = 17-19. Mice aged 10-15 weeks. (F) Fold change in CD8^+^ T cells across various organs in PD-1 KO mice relative to WT mice. Data pooled from three independent experiments, n = 2-3 for lung and kidney, n = 6-10 for spleen, liver, and SMG. FC = 1 indicated by dotted line. (G) PD-1 expression on CD8^+^ T cells across various organs in C57BL/6 WT mice. Data pooled from two independent experiments, n = 4 for all organs, except lungs where n = 2. Data are represented as mean ± SEM. Statistical analysis for B and C utilized a paired t test. Statistical analysis for D and E utilized a two-way Anova. ns P > 0.05, * P ≤ 0.05, ** P ≤ 0.01, *** P ≤ 0.001.

We next examined whether aging influences this phenotype. To do this, PD-1^-/-^ mice and littermate control salivary gland lymphocyte composition was analyzed at various ages from weanling to nearly two years of age **(Supplemental Figure 1G)**. The frequency of CD8^+^ T cells in the salivary glands of weanling mice, as young as 4 weeks of age, are elevated in PD-1 deficient mice as compared to age-matched B6 controls. Additionally, while the frequency of CD8^+^ T cells in the SMG of wildtype mice increases with age, the frequency of CD8^+^ T cells is consistently higher in PD-1^-/-^ mice. In conclusion, PD-1 deficiency significantly impacts the SMG lymphocyte composition, elevating CD8^+^ T cell frequencies in a gender-independent and age-consistent manner, highlighting the role of PD-1 in salivary gland immune regulation.

### Fold change in CD8^+^ T cells is most pronounced in the SMG of PD-1^-/-^ animals

Mucosal tissues are diverse in structure and function and consequently are home to unique resident lymphocyte subsets. We investigated whether the increased number and frequency of CD8^+^ T cells in PD-1^-/-^ mice were confined to the SMG. This was accomplished by comparing wildtype B6 and PD-1^-/-^ lymphocyte composition across various organs of 8–12-week-old mice. No noticeable differences were observed in secondary lymphoid organs such as lymph nodes (not shown) and spleen **(Figure 1F)**. The lung similarly displayed no notable increase in CD8^+^ T cells in PD-1^-/-^ mice **(Figure 1F)**. In contrast, CD8^+^ T cell frequency in the liver and kidneys of PD-1^-/-^ animals was higher than in littermate wildtype controls. We find that there are approximately 3-fold more CD8^+^ T cells in the liver and kidney of PD-1^-/-^ as compared to wildtype animals **(Figure 1F)**. However, the fold change in CD8^+^ T cells was notably most robust in the SMG. Remarkably, there are >10-fold more CD8^+^ T cells in the SMG of PD-1^-/-^ mice as compared to wildtype controls **(Figure 1F).**

It is well known that PD-1 is poorly expressed on lymphocytes from secondary lymphoid organs^12^ including the spleen **(Figure 1G).** In contrast, we find that PD-1 is highly expressed on CD8^+^ T cells from the SMG of naïve B6 mice **(Figure 1G)**, which likely contributes to susceptibility of the SMG to aberrant expansion of CD8^+^ T cells during PD-1 deficiency. Notably, the CD8^+^ T cells of the kidney also express relatively high levels of PD-1 at baseline. Overall, this suggests that high levels of PD-1 expression on CD8^+^ T cells in the naïve SMG is essential in maintaining their homeostasis.

### NK cells regulate CD8^+^ T cells in naïve SMG

We reasoned that the CD8^+^ T cell population is likely to be regulated by a well-represented cell subset in the salivary glands. NK cells, which are the main lymphocyte population in the SMG representing >50% of the CD45^+^ lymphocytes in naïve animals **(Figure 1A and 2A)**, fulfill this requirement. To determine if NK cells control CD8^+^ T cell homeostasis in the SMG, we utilized both depletion and genetic approaches. Because we observed an elevated number of CD8^+^ T cells in the SMG of PD-1^-/-^ mice as young as 4 weeks of age **(Supplemental Figure 1G)**, we reasoned that the regulation of CD8^+^ T cells in the SMG is likely to begin at a neonate age. Therefore, we performed NK cell depletion experiments in weanling mice. To do this, 3-4 week old mice were treated with 200 µg of anti-NK1.1 mAb every 5 days for 3 weeks **(Figure 2B)**. This treatment led to a successful depletion of NK cells in all the organs tested, including the SMG **(Supplemental Figure 2A)**. Spleens and SMG of NK cell-depleted 6-7-week-old mice were then characterized by flow cytometry **(Figure 2C and 2D)**. NK cell depletion led to a significant increase in frequency of CD8^+^ T cells by nearly 7-fold. Importantly, NK cell absence does not significantly change the frequency of CD4^+^ T cells. Notably, this phenotype observed in NK cell depleted animals recapitulated the phenotype observed in PD-1^-/-^ mice **(Figure 1B)**.

**Figure 2:**
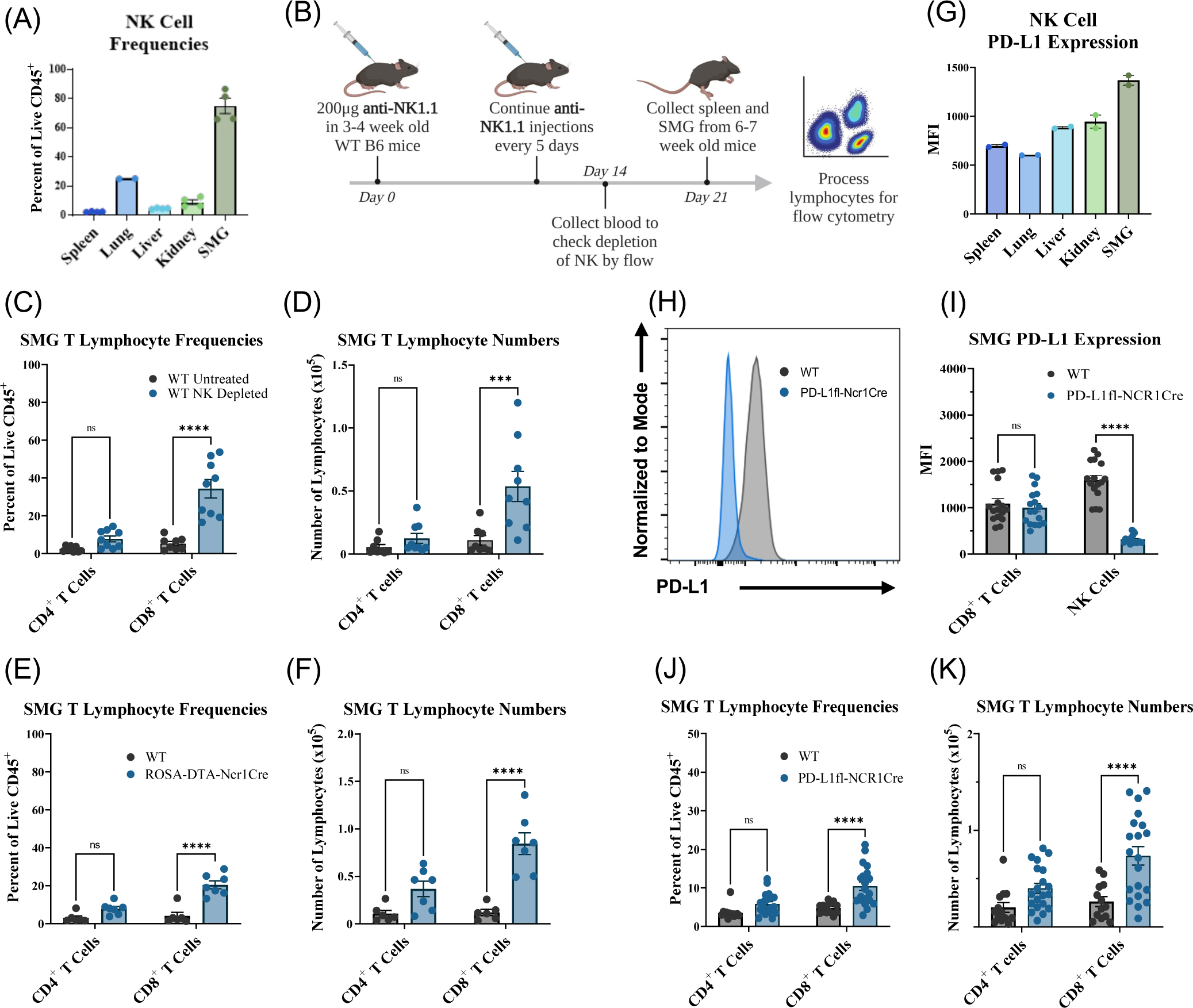
Both NK cell deficiency and PD-L1 deletion on NK cells lead to specific expansion of CD8^+^ T cells in the SMG. (A) Frequency of NK cells across various organs in C57BL/6 WT mice. Data pooled from two independent experiments, n = 4 for all organs, except lungs where n = 2. (B) NK depletion strategy. (C) Frequency and (D) number of CD8^+^ and CD4^+^ T lymphocytes in the SMG of C57BL/6 WT mice after depletion via anti-NK1.1 mAb treatment. Data pooled from three independent experiments, NK depleted (blue), WT controls (grey), n = 9. (E) Frequency and (F) number of CD8^+^ and CD4^+^ T lymphocytes in the SMG of ROSA-DTA-Ncr1Cre mice (blue) and littermate controls (grey). Data from one experiment, n = 6-7. (G) PD-L1 expression on NK cells across various organs in C57BL/6 WT mice. Data representative of several experiments for spleen and SMG, and of one experiment for other organs. (H) Representative histogram validating PD-L1 deletion on NK cells in PD-L1fl-NCR1Cre mice (blue) compared to littermate controls (grey). (I) Expression of PD-L1 on SMG CD8^+^ T and NK cells in PD-L1fl-Ncr1Cre mice (blue) compared to littermate controls (grey). (J) Frequency and (K) number of CD8^+^ and CD4^+^ T cells in PD-L1fl-Ncr1Cre SMG (blue) compared to littermate controls (grey). Data pooled from four independent experiments, n = 13-20. Data are represented as mean ± SEM. Statistical analysis for C-F and I-K utilized a two-way Anova. ns P > 0.05, * P ≤ 0.05, ** P ≤ 0.01, *** P ≤ 0.001.

To complement the depletion approach, we also utilized a genetic approach. This was accomplished by crossing Ncr1Cre mice to ROSA-DTA mice^23^. The ROSA promoter drives expression of diphtheria toxin in the Cre-expressing cells resulting in NK cell ablation^23,24^. In agreement with the depletion experiment, NK cell deficient mice exhibit an increase in the number and frequency of CD8^+^ T cells, but not CD4^+^ T cells in the SMG **(Figure 2E and 2F)**. Altogether these data demonstrate that NK cells regulate CD8^+^ T cell expansion in the salivary glands.

### NK cells regulate CD8^+^ T cells in naïve SMG in part via PD-L1

We next investigated how NK cells regulate CD8^+^ T cell number in the salivary glands. PD-1 has two ligands, PD-L1 and PD-L2^25,26^. PD-L1 is widely expressed by many different hematopoietic and nonhematopoietic cells, while PD-L2 expression is much more restricted and predominantly found on DCs, macrophages, bone marrow–derived mast cells, and B cell populations^27,28^. We find that NK cells of the SMG express high levels of PD-L1 **(Figure 2G)**. These findings combined with the higher PD-1 expression on the expanded CD8^+^ T cells in NK depleted mice **(Supplemental Figure 2B)** further support the potential role of the PD-1/PD-L1 axis between NK and CD8^+^ T cells. We therefore hypothesized that PD-L1 on NK cells controls CD8^+^ T cell expansion in the SMG. To test this hypothesis, we crossed PD-L1 floxed mice^29^ to Ncr1Cre mice. In the resultant mouse line, the only cells lacking PD-L1 are NK cells **(Figure 2H and 2I)**. Of note, the frequency and number of NK cells in the SMG are not altered by PD-L1 deficiency **(Supplemental Figure 2C and 2D)**. Similar to PD-1^-/-^ mice, NK cell-depleted animals, and NK cell-deficient animals, we found that mice lacking PD-L1 on NK cells display a specific increase in CD8^+^ T cells of the SMG, but not CD4^+^ T cells **(Figure 2J and 2K)**. Although not as pronounced as the PD-1 deficient phenotype, these results indicate that PD-L1 on NK cells is responsible for regulating CD8^+^ T cells in the SMG. Importantly, the expanded CD8^+^ T cells express high levels of PD-1 in both the NK cell depleted animals **(Supplemental Figure 2B)** and mice lacking PD-L1 on NK cells **(Supplemental Figure 2E)**. Taken together, these results demonstrate that salivary gland NK cells regulate CD8^+^ T cell expansion in this organ via the PD-1/PD-L1 axis.

### Multimodal single-cell sequencing reveals the heterogeneity of T cells in the mouse SMG

We next examined the nature of the CD8^+^ T cells that accumulate in the salivary glands of PD-1 deficient animals using multimodal single-cell RNA sequencing. CD45^+^ CD3^+^ T cells from the SMG of 12-week-old PD-1^-/-^ mice (n = 6) and wildtype controls (n = 18) were sorted. RNA expression, TCRαβ paired clones and expression of 128 surface proteins **(Supplemental Table 1)** were measured in each single cell in collaboration with ImmGenT in accordance with their standardized protocol^30^. T cells from PD-1^-/-^ individual mice were profiled separately, whereas CD45^+^ CD3^+^ lymphocytes from WT mice were pooled due to low frequency and number (**Figures 1B and 1C**). Females and males were analyzed for both genotypes separately **(Supplemental Table 2)**. For each sample, 300-1000 single cells passed quality control for surface protein expression and RNA expression **(see Methods and Supplemental Figure 4A)**, totaling 5,364 cells post-filtering. Productive TCRα and β sequences were available for 4,646 (87%) of the T cells analyzed.

Unsupervised clustering revealed distinct populations of T cells in the mouse SMG as shown in the Uniform Manifold Approximation and Projection (UMAP) based on RNA data **(Figure 3A)**. These T cell clusters were distinguishable by the expression of key transcripts **(Figure 3B-D)**, and surface expression of key markers **(Supplemental Figure 3)**. Module scores were procured using expression of several key genes to identify the annotations for each cluster **(see Methods and Figure 3C)**. A UMAP from the CITE-seq (Cellular Indexing of Transcriptomes and Epitopes) data on surface protein expression was also generated and overlaid with the clusters generated from the RNA-seq data **(Supplemental Figure 4B)**, showing that the majority of SMG T cells also cluster by CITE-seq protein expression. Differentially expressed CITE-seq surface proteins for each cluster were identified **(Supplemental Figure 4C)**, confirming surface expression of key proteins to identify each cluster.

**Figure 3:**
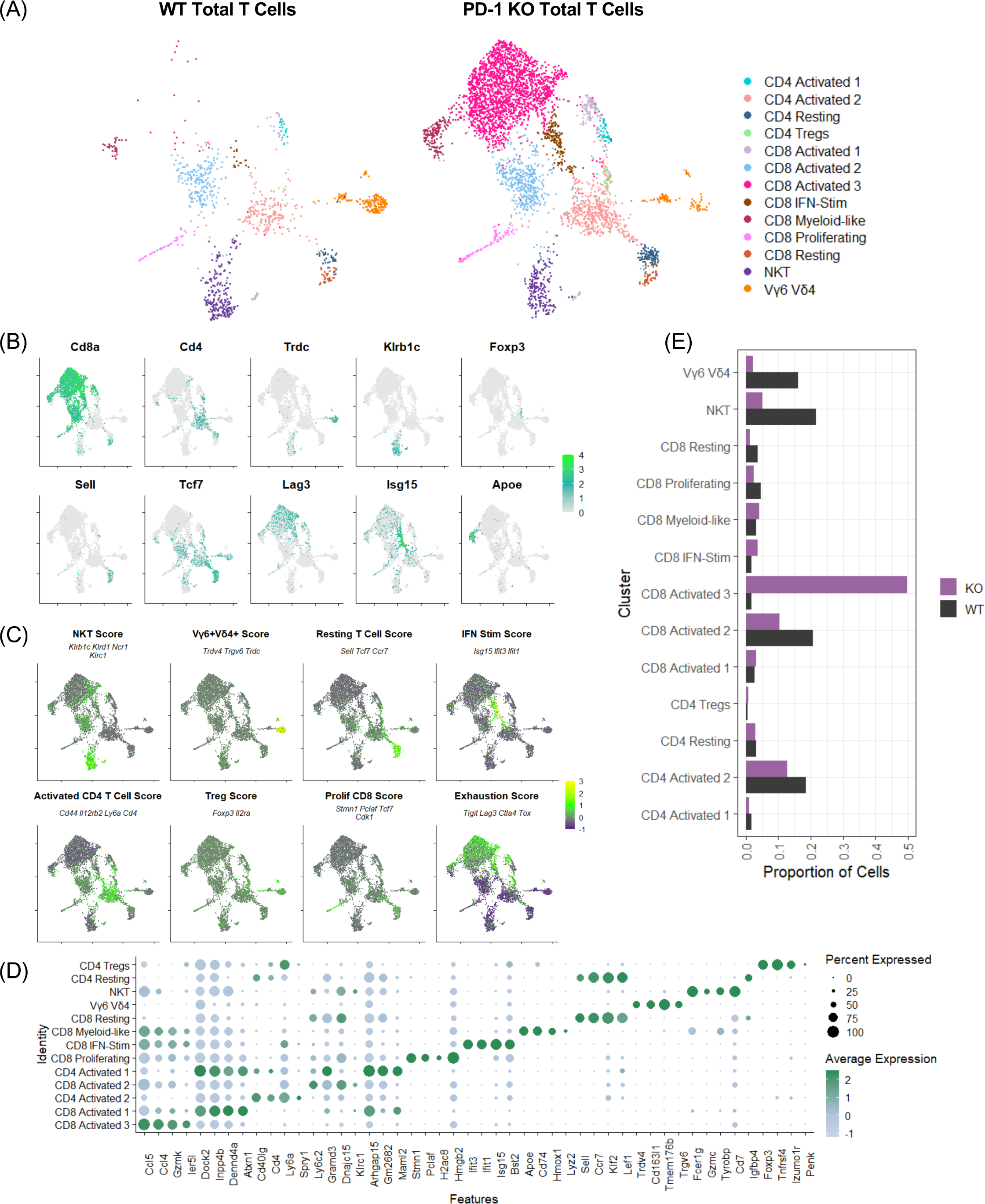
Multimodal single-cell RNA sequencing reveals distinct subsets of T cells in the mouse SMG. Sorted CD45^+^ CD3^+^ T cells from WT and PD-1 KO SMG were subjected to multimodal scRNA-seq (RNA, TCR, CITE-seq). WT samples consisted of 9 male or 9 female SMG pooled together. For PD-1 KO samples, 3 males and 3 females were processed and sequenced individually. (A) Uniform manifold approximation and projection (UMAP) of RNA-seq data from WT control and PD-1 KO mouse SMGs with semi-supervised cluster annotation (see Methods). (B) Feature plots showing log-normalized expression of key genes used to identify clusters. (C) Module score of key genes used to identify clusters (see Methods). (D) Dotplot of enriched genes in each cluster. The dot size represents the proportion of cells in a cluster that express a given feature, while the color indicates the average expression level of that feature across all cells in the cluster (with green representing high expression and light blue indicating low expression). (E) Bar graph representing the proportion of cells per cluster from WT and KO groups.

These identified clusters of SMG T cell subpopulations exhibit differential representation patterns between the WT and PD-1 deficient SMG **(Figure 3E)**. Most notably, the cluster consisting of CD8^+^ T cells with a highly activated phenotype (CD8 Activated 3), was almost exclusively seen in the PD-1 deficient samples, with a 30-fold increase in proportion of cells sequenced between WT and KO samples. Additionally, the interferon stimulated CD8^+^ T cell cluster was enriched within the PD-1 KO samples (CD8 IFN-Stim, 2-fold). In contrast, other subpopulations made up a smaller proportion of the PD-1 deficient T cell population compared to the WT population, including activated CD4^+^ T cells and innate-like T cells (Vγ6Vδ4 and NKT cells) **(Figure 3E)**. SMG from PD-1 deficient animals also harbor decreased proportions of resting CD8^+^ T cells (CD8 Resting, 3-fold decrease) and highly proliferative CD8^+^ T cells (CD8 Proliferating, 2-fold decrease). Taken together, these results highlight the diversity of the T cell subsets present in the naïve mouse SMG and uncover the subpopulations of CD8^+^ T cells that expand in the SMG of PD-1 deficient mice.

### CD8^+^ T cells in the PD-1 deficient SMG display a highly activated transcriptional phenotype

To determine how CD8^+^ T cells in PD-1 deficient SMG differ transcriptionally from wildtype controls, CD8^+^ T cells were isolated from the scRNA-seq dataset. A large number of genes were differentially expressed in PD-1 deficient CD8^+^ T cells **(Supplemental Figure 4D)**, with 12 genes upregulated and 15 genes downregulated (log_2_FC cutoff of 0.5, and a stringent P-value Cutoff of 10e-32). Some of the genes enriched in PD-1 deficient CD8^+^ T cells included proinflammatory and exhaustion genes such as *Ccl5, GzmK, Ier5l, Nr4a2, Rgs16, Tigit, Lag3,* and *Nkg7*.

Because the CD8 Activated 3 cluster is highly enriched within the PD-1 deficient SMG samples, we used our dataset to determine key transcripts associated with this cluster. Differential expression analysis revealed that many of the enriched genes in this cluster **(Figure 4A)** overlap with the markers of heightened activation and exhaustion noted in **Supplemental Figure 4D**. This indicated that the CD8 Activated 3 cluster is driving the phenotype observed in PD-1 KO samples. For instance, the proinflammatory effector molecules *Ccl5* and *GzmK,* are strongly enriched within the CD8 Activated 3 cell cluster **(Figure 4B)**.

**Figure 4:**
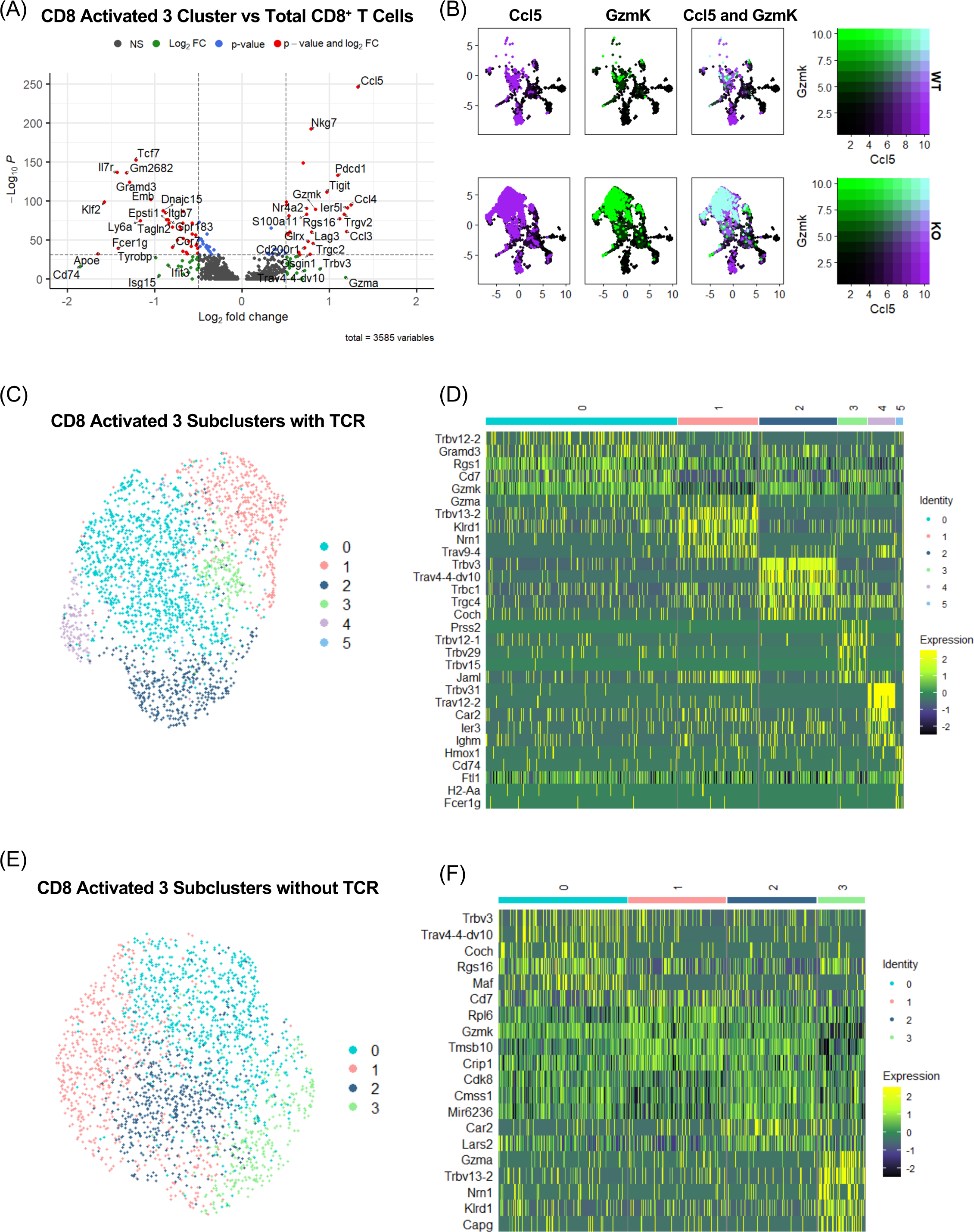
Expanded CD8^+^ T cells in PD-1 KO SMG display a unique transcriptional phenotype. (A) Volcano plot of differentially expressed genes in CD8 Activated 3 cluster compared to total CD8^+^ T cells. P-value cutoff = 10e-32, log_2_FC cutoff = 0.5. Genes highlighted in red meet both the threshold for p-value and log_2_FC, blue only p-value, green only log_2_FC, and grey neither. (B) *Ccl5* and *GzmK* co-expression across all T cells sequenced. (C) scRNA-seq subclusters within CD8 Activated 3 cluster without removal of TCR chains. (D) Heatmap of differentially expressed genes via RNA-seq across subclusters generated within the CD8 Activated 3 population. (E) scRNA-seq subclusters within CD8 Activated 3 cluster after TCR chains were removed from the Variable Features list used for generating the UMAP. (F) Heatmap of differentially expressed genes via RNA-seq across new subclusters generated without TCR chain bias.

To determine the clinical relevance of these highly activated CD8^+^ T cells, we compared them to CD8^+^ T cells associated with Sjogren’s Syndrome in human patients^31,32^. In Sjogren’s Syndrome patients, pathogenic resident GzmK^+^ CD8^+^ T cells expand and correlate with disease severity^32^. Importantly, of 50 genes differentially upregulated in those cytotoxic T cells, 17 are also upregulated in the murine CD8 Activated 3 cluster of interest (**Supplemental Figure 5**). In contrast, there were no more than 6 overlapping genes between the human pathogenic GzmK^+^ CD8^+^ T cells and other clusters we identified (log_2_FC cutoff = 0.25) (**Supplemental Table 3**). These results indicate that the SMG CD8 Activated 3 cluster of PD-1 deficient mice corresponds to the human pathogenic GzmK^+^ CD8^+^ T cell subset described in human patients with primary Sjogren’s Syndrome.

Upon closer inspection of the CD8 Activated 3 cluster, we identified several unique subclusters that clustered by specific TCR α and β chains **(Figure 4C and 4D)**. Therefore, we reran the clustering by omitting the TCR chains from the top variable features used to create the UMAP **(Figure 4E)**. We can still detect TCR chain expression within these unbiased clusters and interestingly, several of the top differentially expressed genes across these subclusters were still specific TCR α and β chains **(Figure 4F)**, indicating that these subclusters contain clonal expansion of specific T cells with similar transcriptional profiles, warranting further detailed analyses of the TCR repertoire.

### Clonal expansion and reduced TCR repertoire diversity is observed in PD-1 deficient SMG

As previously mentioned, in addition to scRNA-seq, we also analyzed T cell receptor variable (TCR-V) sequence (two-chains) at the single-cell level, using IMGT HighVQuest and Immunarch. We identified the top-expanded clonotypes present in our sequenced T cells and found that they predominantly occupy the CD8 Activated 3 cluster enriched in the PD-1 deficient samples **(Figure 5A)**, indicating that clonally expanded CD8^+^ T cells share a transcriptionally similar phenotype. Overall, in comparison to WT controls, the TCR repertoire of PD-1 deficient SMG T cells was less diverse as demonstrated by clonal diversity estimated using Chao1 **(Figure 5B)**, increased repertoire space occupied by a smaller number of specific clones **(Figure 5C)**, and increased abundance of a handful of specific clonotypes in PD-1^-/-^ samples **(Figure 5D)**. In addition to clustering within the CD8 Activated 3 population, some clones are also represented within the CD8 Proliferating, CD8 IFN-Stim, CD8 Myeloid-like, CD8 Activated 1, and CD8 Activated 2 clusters, suggesting that these clones may be precursors and take on a highly activated phenotype following expansion **(Figure 5A and 5E)**. Although some of the most abundant clones only appeared in one mouse, one clone (TRAV9-4.TRAJ31.CAVSANSNNRIFF.TRBV13-2.TRBJ2-7.CASGDREEQYF) was enriched in two different mice (**Figure 5F)**. Importantly, this clone was enriched within the same cluster, suggesting parallel transcriptional pathways following activation.

**Figure 5:**
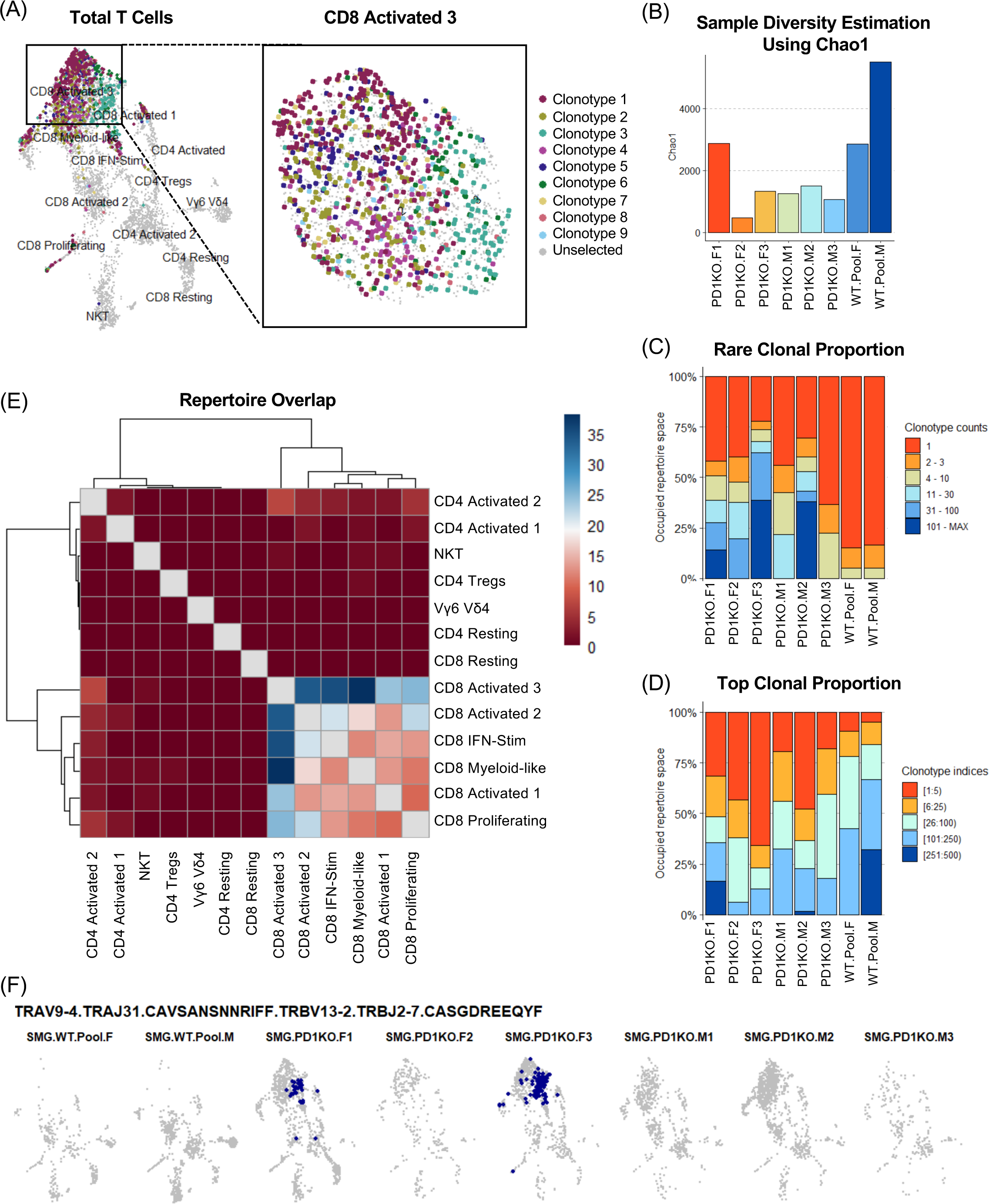
Reduced TCR repertoire diversity in PD-1^-/-^ SMG T cells supports semi-diverse clonal expansion. TCR-seq data was analyzed using IMGT HighVQuest and Immunarch. (A) UMAP overlaid with the most enriched clonotypes. (B) Sample diversity as estimated using Chao1. (C) Proportion of repertoire space occupied by rare clones with specific counts across samples. Rare clones observed only once in a sample are labeled in orange, and clones observed more frequently are shown in blue. (D) Proportion of top clones occupying the repertoire space with the top 5 most abundant clones in each sample labeled in orange. (E) Using Immunarch, shared clonotypes between Seurat clusters were identified. Greater repertoire overlap between two clusters is indicated by dark blue. Comparisons of a cluster to itself were not calculated, and thus are greyed out along the diagonal. Hierarchical trees represent which clusters are most clonally similar. (F) UMAP overlaid with the third most abundant clone, which appears to be clonally expanded in both PD-1 KO female samples 1 and 3.

### TCRdist3 reveals T cell clones with predicted overlapping antigen specificity

After determining the top clones enriched in PD-1 deficient SMG, we sought to determine if any of these clones could have shared antigen specificity. To do so, we enabled the use of TCRdist3^33,34^, which utilizes *in silico* prediction algorithms to identify neighborhoods of TCRs with similar biochemical structure that could theoretically recognize the same antigen. Notably, utilizing the gene-segment pairing plots featured in TCRdist3, we visualized enrichment of specific gene pairings that occupy large proportions of the repertoire space within PD-1 KO SMG samples as compared to the WT SMG repertoire **(Figure 6A and 6B, and Supplemental Figure 6)**.

**Figure 6:**
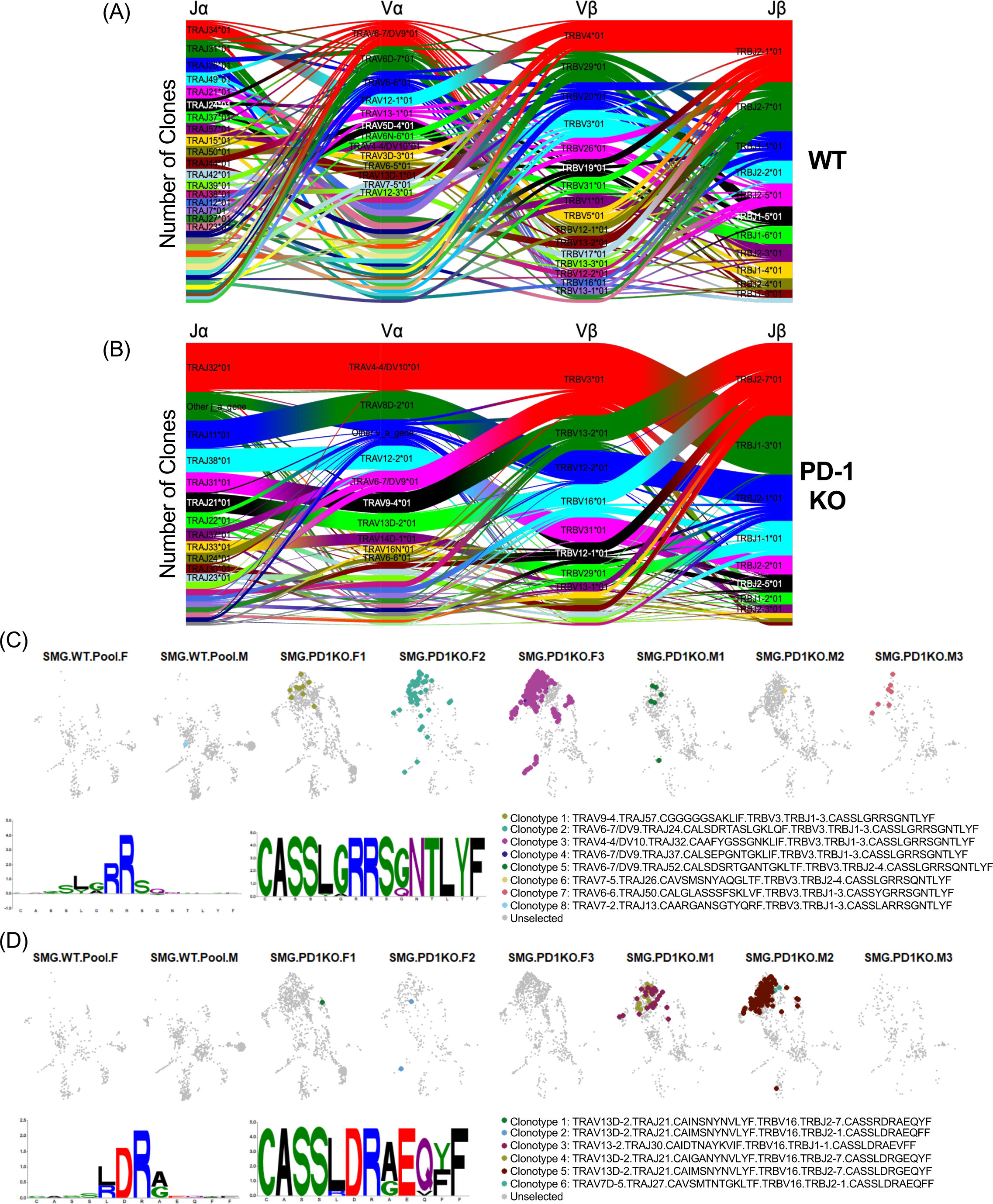
Predicted overlap of antigen recognition in expanded clones via TCRdist3. (A) V gene segment usage and gene pairing landscapes of the top 100 most abundant TCR clones within WT (A) and PD-1 KO (B). Genes are ordered and colored by frequency within the repertoire, with red being the most frequent, then green, blue, cyan, magenta, etc. where the width of the curved connections are proportional to the number of clones in the gene pairing. (C) UMAP overlaid with clones from the most abundant TCRdist3 identified neighborhood and motif analysis of the highlighted quasi-public neighborhood. (D) UMAP overlay and motif analysis of a second identified quasi-public neighborhood.

Moreover, using TCRdist3, we were able to identify several neighborhoods of TCR clonotypes based on their sequences, predicted high biochemical similarity, and TCR distance measures. We identified several quasi-public TCR neighborhoods present in two or more biological replicates and characterized the associated CDR3β motifs **(Figures 6C and 6D)**. For example, the most abundant clone (TRAV4-4/DV10.TRAJ32.CAAFYGSSGNKLIF.TRBV3.TRBJ1-3.CASSLGRRSGNTLYF) was present in 307 T cells in PD-1 KO SMG Female 3, and TCRdist3 analysis identified seven neighboring clones with similarly predicted biochemical structure and antigen specificity present in other mice. Remarkably, these neighboring clones are present in all biological replicates of PD-1 KO SMG T cells, and one of the WT samples, albeit at a low frequency, suggesting that this expanded clone is normally present at a low frequency in the WT SMG but is unleashed when PD-1 is lost **(Figure 6C)**. These clones utilize a shared CDR3β motif CASS(L/Y)(G/A)RRS(G/Q)NTLYF, with varying Vα usage. Markedly, this quasi-public neighborhood shares a similar transcriptional profile and primarily clusters within the CD8 Activated 3 cluster of interest in the PD-1 deficient samples. Notably, this neighborhood stands out in the V gene segment usage and pairing landscape, as seen represented in large part by the thick ribbons connecting TRBV3 to TRAV4-4/DV10 and TRAV6-7/DV9 **(Figure 6B)**. A second quasi-public TCR neighborhood was identified with the CDR3β motif CASS(L/R)DR(A/G)E(Q/Y)(Y/F)F, and occurs across multiple biological replicates of PD-1 KO SMG T cells **(Figure 6D)**. Importantly, these neighboring clones are also localized within the CD8 Activated 3 cluster that is highly enriched within the PD-1 deficient SMG. Taken together, these results indicate that highly activated quasi-clonal CD8^+^ T cells are regulated via PD-1 in this organ.

## Discussion

Our findings highlight that PD-1 deficiency on the otherwise healthy C57BL/6 background can develop immune CD8^+^ T cell infiltrates of the SMG, even at a young age and without prior challenge with infection or tumor. Remarkably, this infiltration is mostly observed in the SMG. Here, we demonstrate that SMG NK cells regulate CD8^+^ T cells via PD-L1, uncovering an unanticipated and distinct regulatory NK mediated mechanism in this organ. Furthermore, our scRNA-seq analysis unveils a highly activated transcriptional phenotype in CD8^+^ T cells within the PD-1 deficient SMG. Particularly interesting is the presence of highly enriched quasi-public clones within the CD8 Activated 3 population, highlighting potential clonal enrichment in response to a shared antigen. Understanding the mechanism of regulation and the identities of these aberrantly expanded CD8^+^ T cells will have significant impact for immune-related adverse event (irAE) treatment and prevention.

Traditional functions of NK cells include direct cytotoxicity through molecules such as perforin and granzyme, enabling them to induce cell death in their targets. NK cells also produce inflammatory cytokines including IFN-γ and TNF-α to activate and recruit other immune cells. Additionally, populations of tissue-resident NK cells have been shown to induce killing of target cells through TRAIL-mediated killing^35,36^. Besides effector functions, recent evidence suggests a regulatory role for NK cells, at least during viral infections. For instance, Schuster and colleagues recently identified a population of tissue-resident memory-like NK cells, which prevent autoimmunity via TRAIL-dependent elimination of CD4^+^ T cells during chronic viral infection^36,37^. Albeit not significant, we observed a mild expansion of the CD4^+^ T cell compartment in NK cell depleted naïve animals. This somewhat corroborates the NK-mediated regulation of CD4^+^ T cells observed by Schuster and colleagues, although the CD4^+^ T cell expansion appears to be more pronounced in infected animals^37^.

In contrast, our data demonstrates a strikingly and significantly increased CD8^+^ T cell compartment in NK cell depleted naïve animals (5-fold increase in number and 7-fold increase in frequency). These findings are consistent with early work from the Pircher group who suggested that NK cells play a regulatory role in restricting the accumulation and development of LCMV specific tissue-resident memory CD8^+^ T cells, although the molecular mechanism was not established^38^. Our data demonstrate that, even in naïve animals, NK cells have alternative regulatory functions that extend beyond their traditional cytotoxic capabilities. Furthermore, our model of PD-L1 deficiency in NK cells suggests that NK cells exhibit different regulation mechanisms of CD4^+^ and CD8^+^ T cells. Regarding the mechanism, our results are consistent with data from Zhou *et al*, who found that liver resident NK cells modulate T cell responses to LCMV infection via the PD-1/PD-L1 axis^39^. However, in our data, NK cell-mediated control of CD8^+^ T cell homeostasis via PD-1 is strongly pronounced in the SMG but significantly less in the liver and kidney, suggesting that the SMG may be a more clinically relevant site to study PD-1 irAEs in the absence of infection. Taken together, our data demonstrate that in naïve animals, NK cells play an immunoregulatory role through their expression of inhibitory ligands, possibly to regulate aberrant CD8^+^ T cells in the SMG via PD-1/PD-L1, to prevent collateral damage and help maintain immune homeostasis.

Another central discovery in our study is the identification and characterization of unleashed CD8^+^ T cells upon PD-1 deficiency. To determine the phenotype of the CD8^+^ T cells expanding in the SMG of PD-1 deficient animals, we employed scRNA-seq, TCR-seq, and CITE-seq. Notably, CD8^+^ T cells with a highly activated signature occupy a drastically greater proportion of the T cells isolated from the SMG of PD-1 deficient mice as compared to wildtype controls. Moreover, PD-1 deficient samples show reduced presence of clusters expressing *Tcf7*, suggesting that these cells expand in the PD-1 deficient SMG and adopt the intensified activated phenotype. Our analyses show that CD8^+^ T cells from naïve PD-1^-/-^ SMG upregulate markers of activation and TCR signaling, including the immediate early response genes *Nr4a2* and *Ier5l*. Furthermore, upregulation of exhaustion markers may indicate that loss of PD-1 results in stimulation and accumulation of CD8^+^ T cells differentiating toward exhaustion. Specifically, PD-1 deficient CD8^+^ T cells upregulate expression of several key exhaustion markers including the inhibitory receptors *Lag3*, *Tigit*, and *Ctla4*, as well as a major transcription factor of exhaustion, *Tox^40,41,^* ^42^. Moreover, RNA expression of *Pdcd1* is highly upregulated in CD8^+^ T cells from the PD-1 deficient SMG, although functional protein expression is absent due to exon 2 deletion. These upregulated inhibitory receptor transcripts indicate a potential negative feedback mechanism whereby PD-1 deficient cells upregulate other inhibitory receptors in an attempt to compensate and regulate heightened activation of the TCR when PD-1 inhibitory signaling is lost. Consequently, lack of PD-1 signaling likely permits continuous expansion and inflammatory effector functions of these highly activated CD8^+^ T cells. Additionally, there was increased expression of *Rgs16*, a regulator of G protein signaling that has been shown to promote antitumor T cell dysfunction and exhaustion^43^. However, our data also show that these unleashed CD8^+^ T cells secrete proinflammatory cytokines and cytotoxic molecules, suggesting that they are not canonically exhausted. We therefore propose that the expanded CD8^+^ T cells in the PD-1 deficient SMG undergo stimulation via TCR activation leading to a heightened activation signature and heading toward aborted exhaustion.

Interestingly, the expanded CD8^+^ T cells of the PD-1^-/-^ SMG also display increased cytokine and chemokine production, including *Ccl5* and *Ccl4*, and they also moderately upregulate the cognate receptor *Ccr5* which has also been seen in other irAEs including myocarditis^44^. Additionally, *Ccl5*, also known as RANTES, has been shown to be significantly upregulated in the saliva of Sjogren’s Syndrome mice. Importantly, the CD8^+^ T cells of interest express high levels of *GzmK*, consistent with highly activated CD8^+^ T cell populations. A recent study from primary Sjögren’s Syndrome patients found a population of CD8^+^ T cells with high expression of *GzmK* that are highly active and cytotoxic^32^. Notably, in this data set, upregulated expression of *GzmK* correlated with disease severity. The authors suggest that the expression of *GzmK* drives the pathogenic inflammation observed in the disease through the production of proinflammatory cytokines as well as the induction of cell death. Importantly, human GzmK^+^ CD8^+^ T cells and murine SMG CD8^+^ T cells share overlapping gene signatures, indicating that they are phenotypically similar. Previous work has also shown that GzmK^+^ CD8^+^ T cells in rheumatoid arthritis patients are potent drivers of inflammatory cytokines^45^. An additional study on T cells associated with inflammaging highlights the accumulation of clonal exhausted-like CD8^+^ T cells expressing high levels of *GzmK*^46^. Overall, we propose that the SMG CD8^+^ T cells expanding in PD-1 deficient animals are the mouse homolog of the GzmK^+^ CD8^+^ T cells identified in Sjögren’s Syndrome patients and possibly other inflammatory conditions including aging.

One recent study on the TCR repertoire following checkpoint blockade in humans found reduction of public CDR3 sequences and expansion of private clones similar to the changes observed in aging or following immunization^47^. In our data set, we also observe expansion of mostly private, not public clones. However, TCRdist3 analysis revealed neighborhoods of quasi-public TCR clonotypes spread across biological replicates. These neighborhoods represent clonotypes with potential shared antigen specificity that may expand in response to the same antigen within the SMG. Notably, the top neighborhood contains enriched clones across 5 out of 6 PD-1 KO biological replicates. Importantly, even for a given epitope (GP33), it has recently been shown that TCR defining individual clones are diverse, highlighting the need for TCR Ag binding tool predictions^48^. This identified neighborhood of clones demonstrates that a common population of CD8^+^ T cells present at low levels and regulated in the WT SMG can expand in PD-1 deficient animals where peripheral tolerance is disrupted. It is well known that negative selection in the thymus is imperfect, and therefore peripheral regulation of self-reactive clones is necessary to maintain tolerance^49,50^. However, loss of these regulatory mechanisms lowers the threshold of activation by self-antigens, resulting in clonal proliferation and autoimmunity. While we observe an expansion of certain TCR clones, additional work will be necessary to determine whether the expanded clones are reacting to a self-antigen.

In summary, our findings challenge the notion that NK cell functions are solely characterized by their capacity to produce cytokines and engage in cytotoxic killing. We provide evidence that NK cells play a crucial role in controlling CD8^+^ T cell expansion in the SMG. Their absence or inability to regulate may contribute to the development of salivary gland dysfunction seen in PD-1 triggered irAEs. Importantly, our data demonstrate that the regulated CD8^+^ T cells have limited TCR diversity and are likely to be the homolog of GzmK^+^ CD8^+^ T autoimmune cells identified in humans with primary Sjögren’s Syndrome. These findings may have important implications for the development of new therapies aimed at preventing or treating salivary gland dysfunction for these patients. By harnessing these immunoregulatory NK cells, we can help protect delicate organs including the salivary glands from irAEs.

## Supporting information

Borys Supplementary figures

## Acknowledgments

This work was supported by the following NIH grants: RO1 AI-046709 (L.B.), RO1 AI-173163 (L.B.), R24 AI-072073 (ImmGen), and F31 DE-032593 (S.M.B.). S.P.R. was supported by a Research Supplement 3R01AI046709-19S1 to promote diversity. S.M.B. was also supported by a Brown University Presidential Award. We would like to thank the ImmGen consortium for their expertise and funding for the single-cell experiments. We would also like to thank Kevin Carlson for assistance with cell sorting, Delia Demers for her expertise in mouse handling, Dania Mallah for assistance with single-cell analysis, and Dr. Eric Vivier for providing NCR1Cre mice.

## Author Contributions

S.M.B. and L.B. conceptualized the study, designed experiments, analyzed data, and wrote the manuscript with input from other authors. S.M.B. performed experiments with mice and prepared samples for single-cell techniques. I.M. prepared the single-cell library for sequencing. S.M.B., I.M., and S.P.R. analyzed the single-cell data with mentorship from D.Z.

## Declaration of Interests

The authors declare no competing interests.

## Inclusion and diversity

We support inclusive, diverse, and equitable conduct of research.

## Data and code availability

Single-cell RNA sequencing data are available at Gene Expression Omnibus (GEO) under accession numbers listed in the key resources table. Any additional information required to reanalyze the data will be shared by the lead contact upon request.

## Materials and Methods

### Mice

C57BL/6 (Strain #000664), ROSA-DTA (Strain #009669)^23^, and PD-L1 floxed (Strain #036255)^29^ mice were purchased from The Jackson Laboratory. NCR1Cre mice were kindly provided by Dr. Vivier^24^. PD-1^-/-^ mice were generated in conjunction with the Brown University Mouse Transgenic and Gene Targeting Facility. Two single-guide RNAs (sgRNAs) were generated to target *Pdcd1* 5’ upstream of exon 2 in the surrounding intron: 64_Pdcd1_sgRNAup1: AGGAGCTTGTAGCTTCTTGT and 65_Pdcd1_sgRNAup2: GAAGCTACAAGCTCCTAGGT. Additionally, two sgRNAs were generated to target 3’ downstream of exon 2 in the surrounding intron: 66_Pdcd1_sgRNAdw1: GGCATCTGAGGATTTCCACA and 67_Pdcd1_sgRNAdw2: GGGCATCTGAGGATTTCCAC. sgRNAs were selected using the CRISPR Guide RNA design tool Benchling (https://www.benchling.com/crispr/). Cas9/plasmid injection was performed on C57BL/6NJ zygotes. Non-homologous end-joining led to a 632 bp deletion of the sequence between sgRNAs resulting in loss of exon 2 of *Pdcd1*. Founders were genotyped via long-range PCR and sequencing at the *Pdcd1* locus. Long-range PCR primers of PD-1^-/-^ mice: 76_Pdcd1_upF1: TCCCACTGACCCTTCAGACAG and 79_Pdcd1_RHdR2: CGTGTCAGGCACTGAAGAGATC. Products were subsequently sequenced with 77_Pdcd1_upF2: AACTAGGCTAGCCAACCAGAAG and 81_Pdcd1_seqF: ACAGTGGCATCTACCTCTGTGG. The resulting mouse line was backcrossed to C57BL/6J (Strain #000664) from The Jackson Laboratory several times. Both age- and sex-matched mice were used for this study, which was carried out in strict accordance with the recommendations in the Guide for the Care and Use of Laboratory Animals, as defined by the National Institutes of Health (D16-00183). Animal protocols were reviewed and approved by the Institutional Animal Care and Use Committee (IACUC) of Brown University. All animals were housed in a centralized and AAALAC-accredited research animal facility that is fully staffed with trained husbandry, technical, and veterinary personnel.

### Isolation of Lymphocytes

Mice were humanely euthanized with isoflurane and cardiac puncture or cervical dislocation prior to organ removal. Spleens were dissociated in 150 mM NH4Cl for 10 minutes, filtered through nylon mesh, and washed twice with 1% PBS-serum. SMGs were processed manually to remove the sublingual glands, parotid glands, and lymph nodes. The isolated SMG was digested in 2 mL digestion solution (DMEM (HyClone) with 16.7 µg/mL Liberase-DL (Sigma-Aldrich) and 0.3 U/mL DNase I (Sigma-Aldrich)) on the heart01.01 program on the GentleMACS and incubated at 37°C for 10 min. This process was repeated for a total of two dissociations and two incubations. After digestions, the SMG samples were filtered through nylon mesh, and washed once with 1% PBS-serum before a Lympholyte-M gradient underlay. Lymphocytes were harvested from the gradient interface and washed once in 1% PBS-serum. Kidneys were cut into small pieces followed by Liberase-DL digestion for 30 min at 37°C, then washed in 1% PBS-serum before purification on a 40%/70% Percoll gradient centrifugation (2500 RPM) at RT. Lymphocytes were harvested from the gradient interface and resuspended in 1% PBS-serum. Livers were perfused with 1% PBS-serum before removal, dissociated with the E.01 program on a GentleMACS in 1% PBS-serum, and filtered through nylon mesh. The liver samples were subsequently washed three times with 1% PBS-serum, then subjected to a 40%/70% Percoll gradient centrifugation at RT. Liver lymphocytes were harvested from the gradient interface and washed once with 1% PBS-serum. Lungs were perfused with 1% PBS-serum prior to removal, dissociated on the lung01.02 program on the GentleMACS, incubated with Liberase-DL for 60 minutes at 37°C, then digested a second time on the lung02.01 program on the GentleMACS. The lung samples were then washed before isolating the lymphocytes with a 40%/70% Percoll gradient centrifugation. Lymphocytes were harvested from the gradient interface and resuspended 1% PBS-serum.

### Flow Cytometry

Single cell suspensions were stained in 1% PBS-serum containing Fc block (2.4G2, produced in house), Brilliant Stain Buffer (BD Biosciences), and fluorochrome-conjugated monoclonal antibodies for 20 minutes on ice in the dark. Dead cells were excluded using Zombie UV (Biolegend), Zombie NIR (Biolegend), or 4’, 6-diamidino-2-phenylindole hydrate (DAPI). Cells were acquired using a four-laser Cytek Aurora Spectral Flow Cytometry System, FACSCelesta (BD Biosciences), or FACSAria III (BD Biosciences) and analyzed using FlowJo (Tree Star Inc.).

### *In Vivo* Treatments

NK cells were depleted from C57BL/6 mice starting at 3-4 weeks of age by intraperitoneal injection of 200 µg of anti-NK1.1 mAb (clone PK136, Bio X Cell) every 5 days for 3 weeks.

### Statistical Analysis of Flow Cytometry Data

Statistical significance was determined using Prism 7.0 (Graph-Pad Software, Inc.). Unpaired two-tailed Student’s t tests were used to compare two individual groups, whereas an ANOVA with multiple comparison test was used when more than two groups were compared. Graphs represent the mean and error bars indicate standard error of the mean (SEM). The number of animals/sample size and total experimental replicates are noted in each figure legend. *p < .05, **p < .01, ***p < .0001, and ****p < .00001.

### Single-cell RNA, TCR and TotalseqC-Sequencing - Sample Preparation

*Flow cytometry and HT antibody staining.* Single cell lymphocyte suspensions were isolated as described above and stained with anti-CD45-APC (Biolegend 103112, clone 30-F11), anti-CD3e-PE (Biolegend 100308, clone 145-2C11), and different TotalSeq-C Anti-Mouse Hashtags (TotalSeq-C Anti-Mouse Hashtags 1-10 Biolegend #155861-155879 **(Supplemental Tables 1 and 2)**, in staining buffer (DMEM, 2% FCS and 10mM HEPES) for 20min at 4°C in the dark. Spleen standard cell suspensions were stained in the same conditions but used anti-CD45-FITC (Biolegend 109806, clone 104) to distinguish it from the other samples once pooled. Each sample was stained with a single hashtag, which allows for the downstream assignment of individual 10x cell barcodes to their source samples^51^. *Cell sorting.* For each sample, 4-15K DAPI^-^ CD45^+^ CD3^+^ T cells were sorted on the FACSAria III and pooled in a single collection tube at 4C. DAPI was added just before the sort. *ImmGenT TotalSeq-C custom mouse panel staining.* The pooled single cell suspension was stained with the ImmGenT TotalSeq-C custom mouse panel, containing 128 antibodies (Biolegend Part no. 900004815) (**Supplemental Table 1**) and FcBlock (Bio X Cell #BE0307). Because 500,000 cells are required for staining, unstained splenocytes were spiked in to reach a total of 500,000 cells. *Second sort.* Cells were subsequently sorted a second time with the addition of DAPI to select for live CD45^+^ CD3^+^ T cells. 20,000 CD45^-^ APC^+^ sample cells and 10,000 CD45^-^ FITC^+^ spleen standard cells were sorted into a single collection tube.

### Single-cell RNA, TCR and TotalseqC-Sequencing - Library Preparation

The collected cells were pooled and split into two technical replicates (independent libraries) for all subsequent steps. Single-cell RNA-seq was performed using the 10x Genomics 5’v2 with Feature Barcoding for Cell Surface Protein and Immune Receptor Mapping following the manufacturer’s instructions (CG000330). After cDNA amplification, the smaller fragments containing the Totalseq-C derived cDNA were purified and saved for Feature Barcode library construction. The larger fragments containing transcript-derived cDNA were saved for TCR and Gene Expression library construction. For both cDNA portions, library size was measured by an Agilent Bioanalyzer 2100 High Sensitivity DNA assay and quantified using a Qubit dsDNA HS Assay kit on a Qubit 4.0 Fluorometer. *TCRαβ amplification and library construction.* TCR cDNA was amplified from the transcript-derived cDNA by nested PCR according to the manufacturer’s protocol. The TCR libraries were then subjected to enzymatic fragmentation, ligation, and indexing with unique Dual Index TT set A (10x part no. 3000431) index sequences, TT-A9 and TT-A10, with an index PCR (8 cycles). *RNA library construction.* Transcript-derived cDNA was processed into the RNA libraries. Following enzymatic fragmentation and size selection, purified cDNA was ligated to an Illumina R2 sequence and indexed with unique Dual Index TT set A index sequences, TT-H6 and TT-H7, with an index PCR (14 cycles). *TotalSeq-C library construction*. Totalseq-C derived cDNA was processed into the Feature Barcode libraries. The cDNA weas indexed with unique Dual Index TN set A (10x part no. 3000510) index sequences, TN-C8 and TN-C9 with an index PCR (8 cycles). *Sequencing of libraries.* The three libraries were pooled together based on molarity with relative proportions 47.5% RNA, 47.5% Feature Barcode, and 5% TCR. This pool was then sequenced on an Illumina Novaseq S2, 100c. The sequencing configuration followed 10x specifications, namely 26 cycles in Read 1, 10 in Index 1, 10 in Index 2, and 90 in Read 2.

### Single-cell RNA, TCR and TotalseqC-sequencing - Data processing

*Count matrices*. Gene and TotalSeqC antibody (surface protein panel and hashtags) counts were obtained by aligning reads to mm10(GRCm38) and the M25(GRCm38.p6) GeneCode annotation (GRCm38.p6) of the mouse genome and the DNA barcode for the TotalSeqC panel (**Supplemental Table 1**) using CellRanger software (v7.1.0) (10X Genomics) with default parameters. Cells were distinguished from droplets with high RNA and TotalSeqC counts using inflection points on total count curve (barcodeRanks function in the DropletUtils package). *Sample demultiplexing*. Sample demultiplexing used the Hashtag counts and the HTODemux from the Seurat package (Seurat v4.1). Doublets (droplets with two hashtags) were excluded and cells were assigned to the max hashtag signal if it had at least 10 counts and more than twice the signal for the second most abundant hashtag. The hashtag count data were also analyzed by t-SNE for a visual check (clear separated clusters for each hashtag). All single cells from the gene count matrix were matched unambiguously to a single hashtag (and therefore their original sample). *Quality control (QC) and batch correction.* Cells matching at least one of the following criteria were excluded: fewer than 500 RNA counts; dead cells with more than 10% of counts mapping to mitochondrial genes; less than 500 TotalSeqC counts; cells positive for 2 isotype controls (non-specific binding of antibodies). One batch experiment was performed and included biological replicates. No significant batch effect was observed after examination of UMAP. *Clustering and dimensionality reduction.* Data from the two technical replicates were merged and normalized using centered log ratio (CLR). The top 2000 variable genes were found using the FindVariableFeatures function in Seurat (v4.2.0) and the variance-stabilizing transformation (VST method). PCA was performed using the top 2000 genes. The top principal components explaining 80% of the total variance were used for two-dimensional reduction using UMAP. Clustering was performed on the same PCs using the FindClusters() function in Seurat. *Differential Expression.* To determine differentially expressed genes, FindMarkers() within Seurat was used. AddModuleScore() was used to visualize aggregated expression of a set of genes. Volcano plots were created using EnhancedVolcano (v1.14.0). *TCRαβ analyses.* TCRαβ contigs for each single cell were obtained by aligning reads to reference genes (refdata-cellranger-vdj-GRCm38-alts-ensembl-7.0.0) using CellRanger software (v7.1.0) with default parameters. Immunarch (v1.0.0) was used to analyze TCRαβ repertoires. TCR gene usage and quasi-public clone neighborhood analyses were performed with TCRdist3^33^ using Python (v3.10.10) and Jupyter notebook (v6.5.4).

## Supplemental Figure Legends

<u>Supplemental Figure 1: Origination and validation of PD-1 KO mice used in study.</u> (A) Map of CRISPR/Cas9 mediated deletion of *Pdcd1* exon 2, created in BioRender. (B) Confirmation of PD-1 deletion on SMG CD8^+^ T cells in the PD-1 KO mice compared to WT and Het littermate controls, n = 4-6. (C) Frequency of CD8^+^ T cells in naïve SMG of WT, in house derived PD-1 KO, and PD-1 KO available from Jackson Laboratories, n = 4. (D) Representative gating strategy of CD45 lymphocytes from naïve WT and PD-1 KO SMG. (E) CD8^+^ T cell frequency and (F) number in WT vs PD-1 KO SMG, colored by sex. Data pooled from 6 independent experiments, n = 17-19. Mice aged 10-15 weeks. (G) SMG CD8^+^ T cell frequency with age in WT vs PD-1 KO SMG. Data pooled from 5 independent experiments, n = 29. Data are represented as mean ± SEM. Statistical analysis for B and C utilized a one-way Anova. Statistical analysis for E and F utilized a paired t test. ns P > 0.05, * P ≤ 0.05, ** P ≤ 0.01, *** P ≤ 0.001.

<u>Supplemental Figure 2: Validation of NK depletion in SMG and expression of PD-1 on expanded CD8^+^ T cells.</u> (A) Frequency of NK cells in the SMG from WT mice treated with anti-NK1.1 depletion antibodies compared to untreated WT mice. (B) PD-1 expression on CD8^+^ T cells following NK depletion. Data for (A) and (B) pooled from three independent experiments, NK depleted (blue), WT controls (grey), n = 9, PD-1 KO reference control (purple), n = 3. (C) Frequency and (D) number of NK cells in SMG of PD-L1fl-Ncr1Cre mice compared to littermate controls. (E) Expression of PD-1 on SMG CD8^+^ T cells of PD-L1fl-Ncr1Cre mice compared to controls. Data for (C), (D), and (E) pooled from four independent experiments, n = 13-20. Data are represented as mean ± SEM. Statistical analyses utilized a paired t test. ns P > 0.05, * P ≤ 0.05, ** P ≤ 0.01, *** P ≤ 0.001.

<u>Supplemental Figure 3: Validation of Seurat cluster annotations using CITE-seq marker gating strategy.</u> Confirmation of innate T cell clusters by surface protein expression of KLRBC-NK1.1 for NKT (A) and TCRGD for Vγ6Vδ4 (B). (C) Cells from clusters that did not include the innate like T cells (NKT and Vγ6Vδ4 clusters) were then subdivided based on CD8A and CD4 surface protein expression. (D) CD44 and CD62L expression on CD4^+^ T cells confirmed resting and activated populations. (E) Surface protein expression of IL2RA.CD25 and RNA expression of Foxp3 on CD4^+^ T cells identified CD4^+^ T regs (F) CD44 and CD62L expression on CD8^+^ T cells validated resting and activated populations.

<u>Supplemental Figure 4: Differential expression of transcriptomes and epitopes across SMG T cells.</u> (A) Number of cells sequenced post filtering in each sample. (B) UMAP based on CITE-seq data, overlaid with RNA-seq clusters. (C) Heatmap of differentially expressed surface proteins across the original Seurat clusters identified via RNA-seq data in Figure 5, with highly expressed markers in yellow and lowly expressed markers in dark blue. (D) Volcano plot of differentially expressed genes in PD-1 KO vs WT CD8^+^ T cells. P-value cutoff = 10e-32, log_2_FC cutoff = 0.5. Genes highlighted in red meet both the threshold for p-value and log_2_FC, blue only p value, green only log_2_FC, and grey neither. (E) Genes enriched within the CD8 Activated 3 cluster compared to total CD8^+^ T cells with log_2_FC > 0.5 were identified for Gene Set Enrichment Analysis using Molecular Signatures Database (MSigDB) derived from the Gene Ontology Biological Process (GO:BP) as a custom source for g:Profiler. The number of enriched genes associated with each gene set are indicated by the bubble size. The precision, which is the proportion of genes in the input list that are annotated to the function for this gene set, is indicated by color where green is more precise while blue is less precise.

<u>Supplemental Figure 5: CD8 Activated 3 cluster from murine SMG mirrors GzmK^+^ CD8^+^ T cells from Sjögren’s Syndrome patients.</u> (A) Differentially expressed genes specific to the CD8 Activated 3 cluster compared to all other SMG T cells were identified with log_2_FC > 0.25. The intersection of genes compared to the top 50 differentially expressed genes specific to the GzmK^+^ CD8^+^ T_M_ cluster identified from human Sjögren’s Syndrome patients are visualized with a Venn diagram. (B) Feature plots showing the expression of 17 overlapping genes found between these two differentially expressed gene sets, with high expression in blue and low expression in grey.

<u>Supplemental Figure 6: Extended clone analysis of WT and PD-1 KO repertoires via TCRdist3.</u> V gene segment usage and gene pairing landscapes of all clones within WT Spleen control (A), WT SMG (B), and PD-1 KO SMG (C).

## Notes

### Competing Interest Statement

The authors have declared no competing interest.

