## Supplementary material for "PD-1 Mediated Regulation of Unique Activated CD8^+^ T Cells by NK Cells in the Submandibular Gland": Borys Supplementary figures

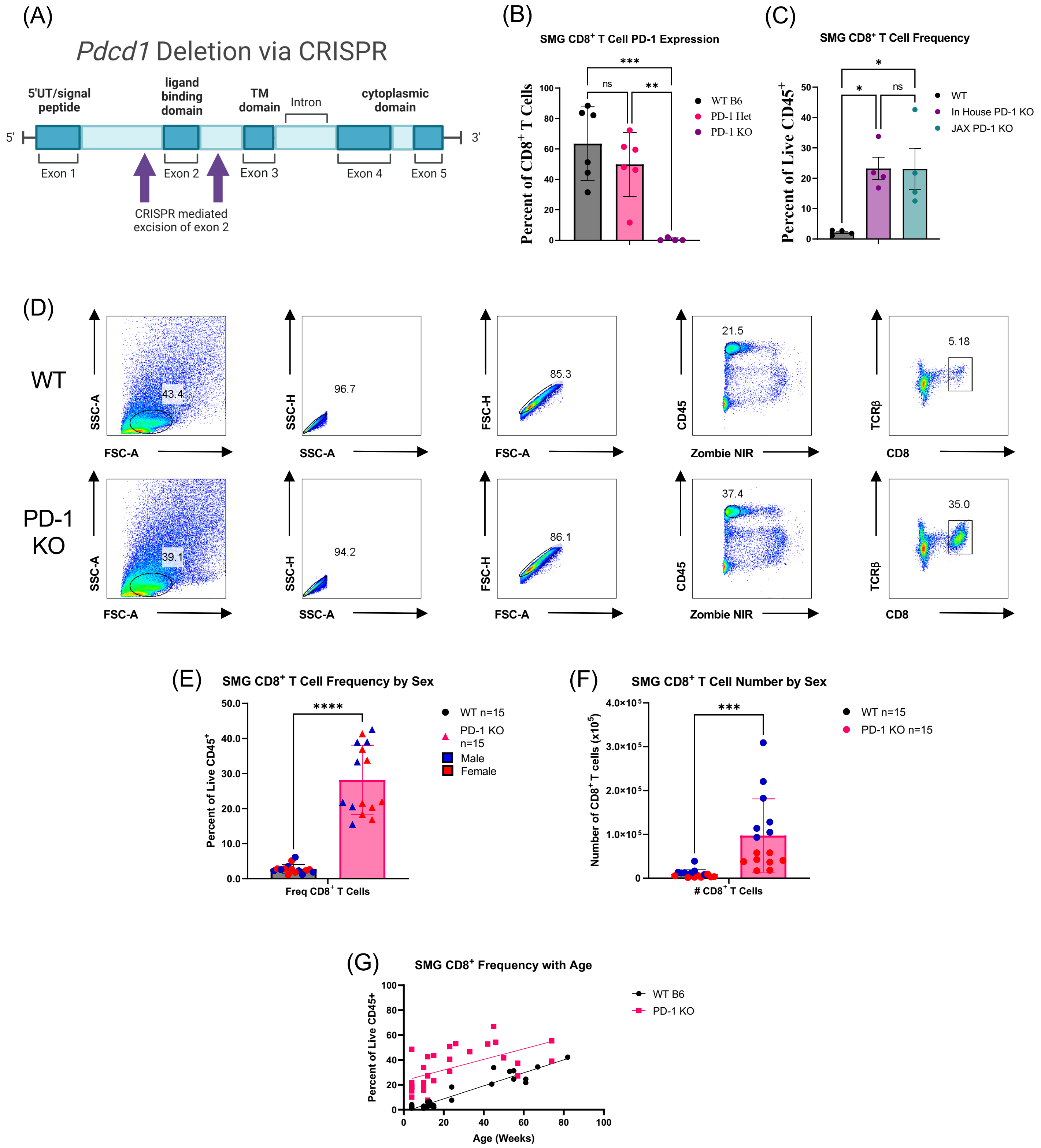


Supplemental Figure 1

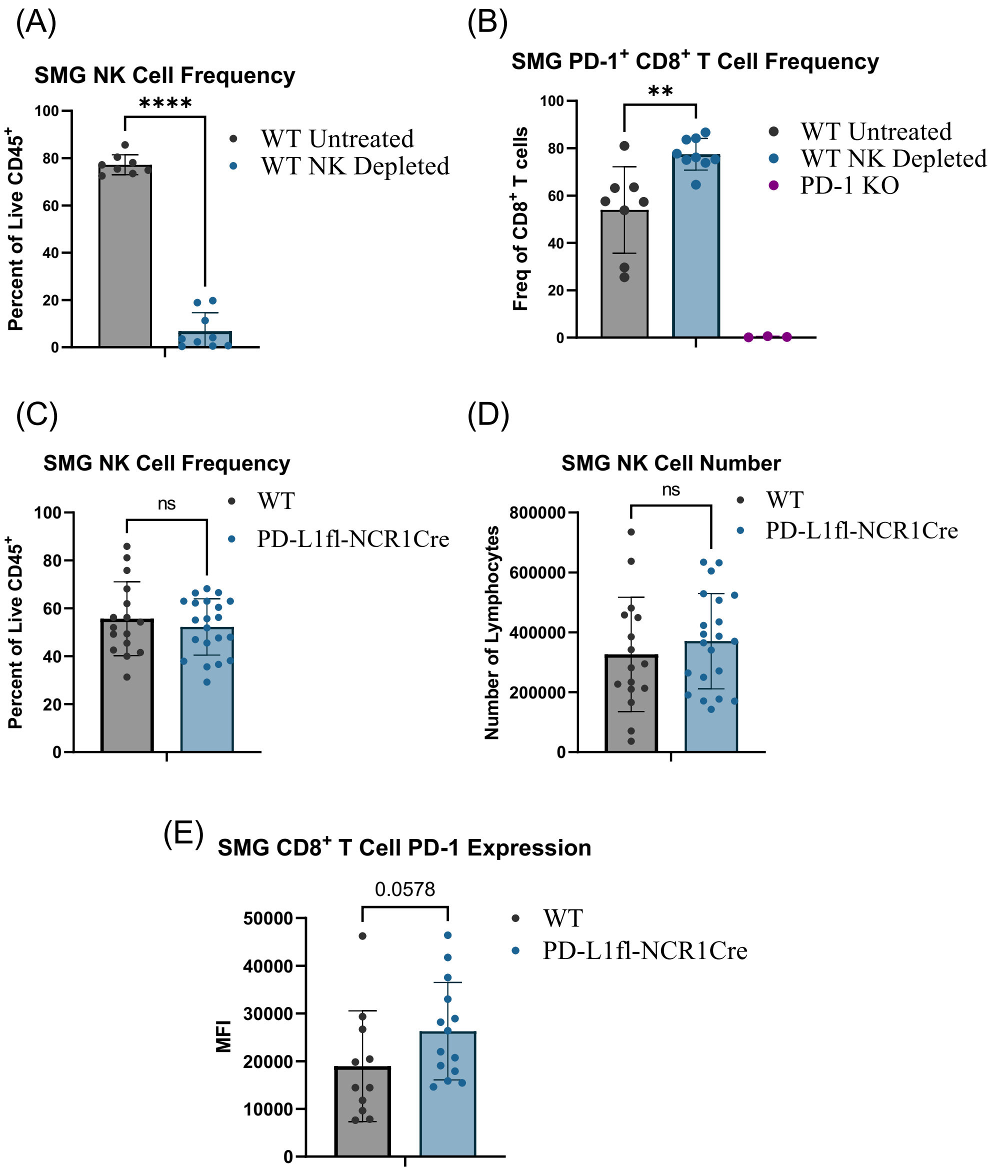
(A)
SMG NK Cell Frequency
****
Percent of Live CD45+
WT Untreated
WT NK Depleted
(B)
SMG PD-1+ CD8+ T Cell Frequency
**
Freq of CD8+ T cells
WT Untreated
WT NK Depleted
PD-1 KO
(C)
SMG NK Cell Frequency
ns
Percent of Live CD45+
WT
PD-L1fl-NCR1Cre
(D)
SMG NK Cell Number
ns
Number of Lymphocytes
WT
PD-L1fl-NCR1Cre
(E)
SMG CD8+ T Cell PD-1 Expression
0.0578
MFI
WT
PD-L1fl-NCR1Cre

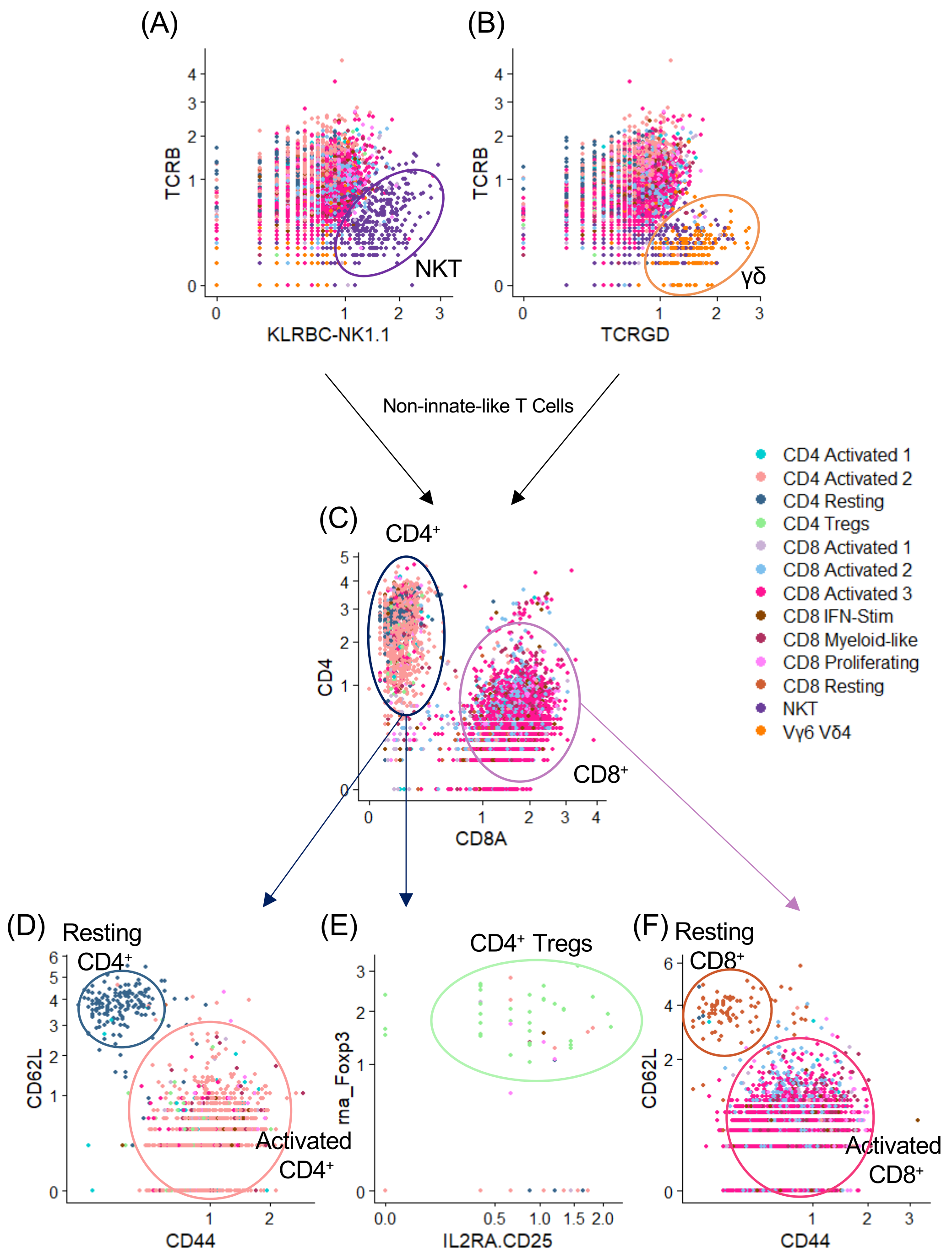
(A)
(B)
TCRB
KLRBC-NK1.1
NKT
TCRGD
γδ
Non-innate-like T Cells
(C)
CD4⁺
CD4
CD8A
CD8⁺
CD4 Activated 1
CD4 Activated 2
CD4 Resting
CD4 Tregs
CD8 Activated 1
CD8 Activated 2
CD8 Activated 3
CD8 IFN-Stim
CD8 Myeloid-like
CD8 Proliferating
CD8 Resting
NKT
Vγ6 Vδ4
(D)
Resting CD4⁺
CD62L
CD44
Activated CD4⁺
(E)
CD4⁺ Tregs
rna_Foxp3
IL2RA.CD25
(F)
Resting CD8⁺
CD62L
CD44
Activated CD8⁺

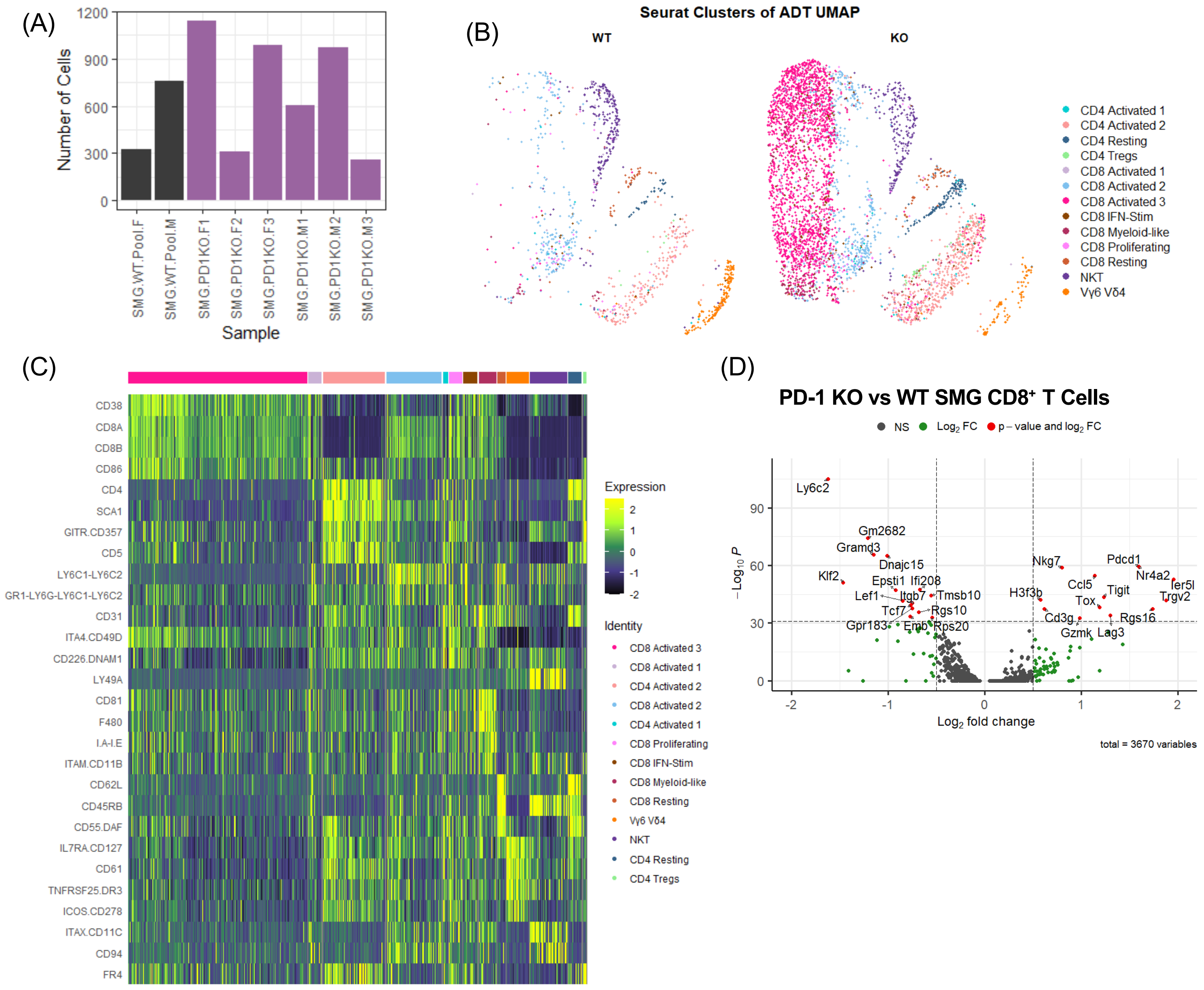


Supplemental Figure 4

(A)

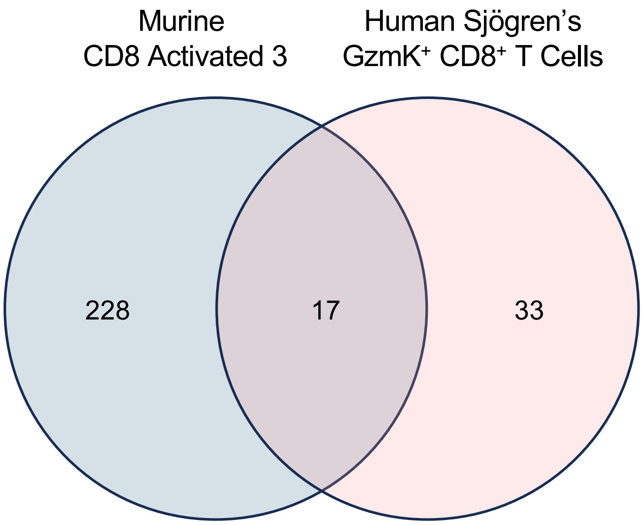
Murine
CD8 Activated 3
Human Sjögren's
GzmK+ CD8+ T Cells
228
17
33

(B)

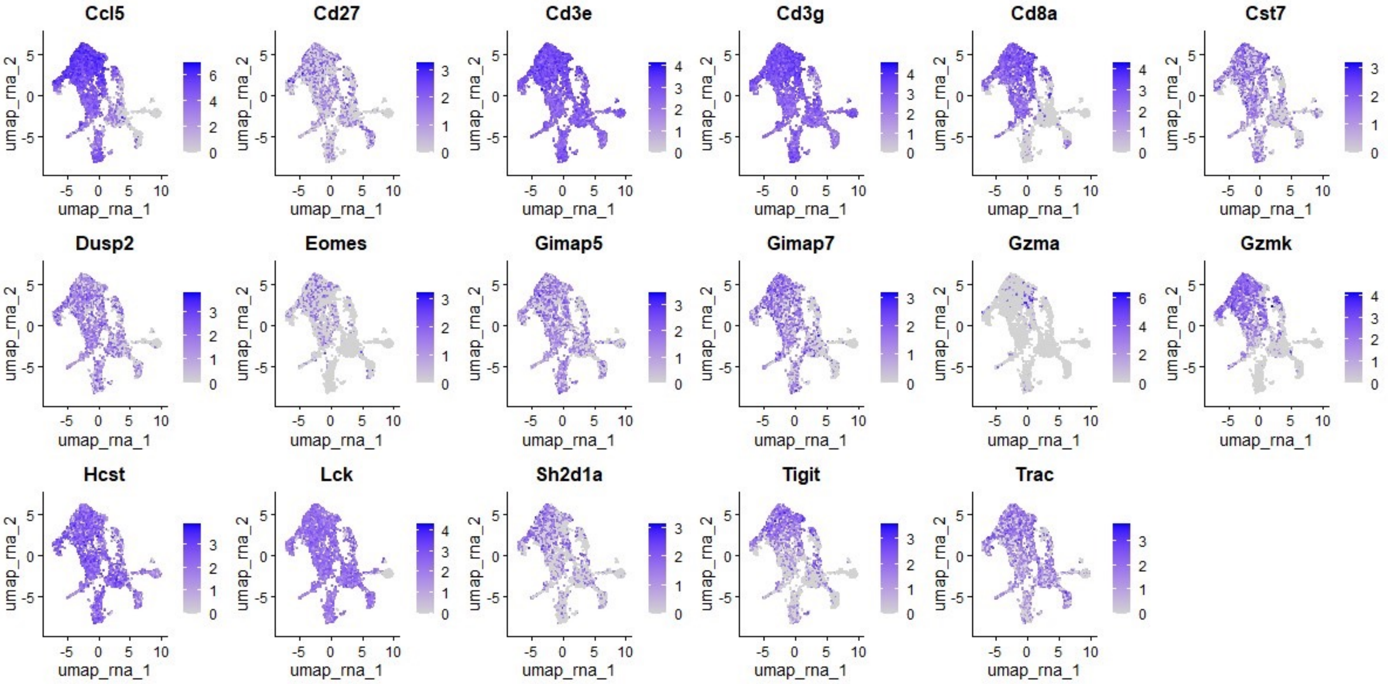
Ccl5
Cd27
Cd3e
Cd3g
Cd8a
Cst7
Dusp2
Eomes
Gimap5
Gimap7
Gzma
Gzmk
Hcst
Lck
Sh2d1a
Tigit
Trac
umap_rna_1
umap_rna_2

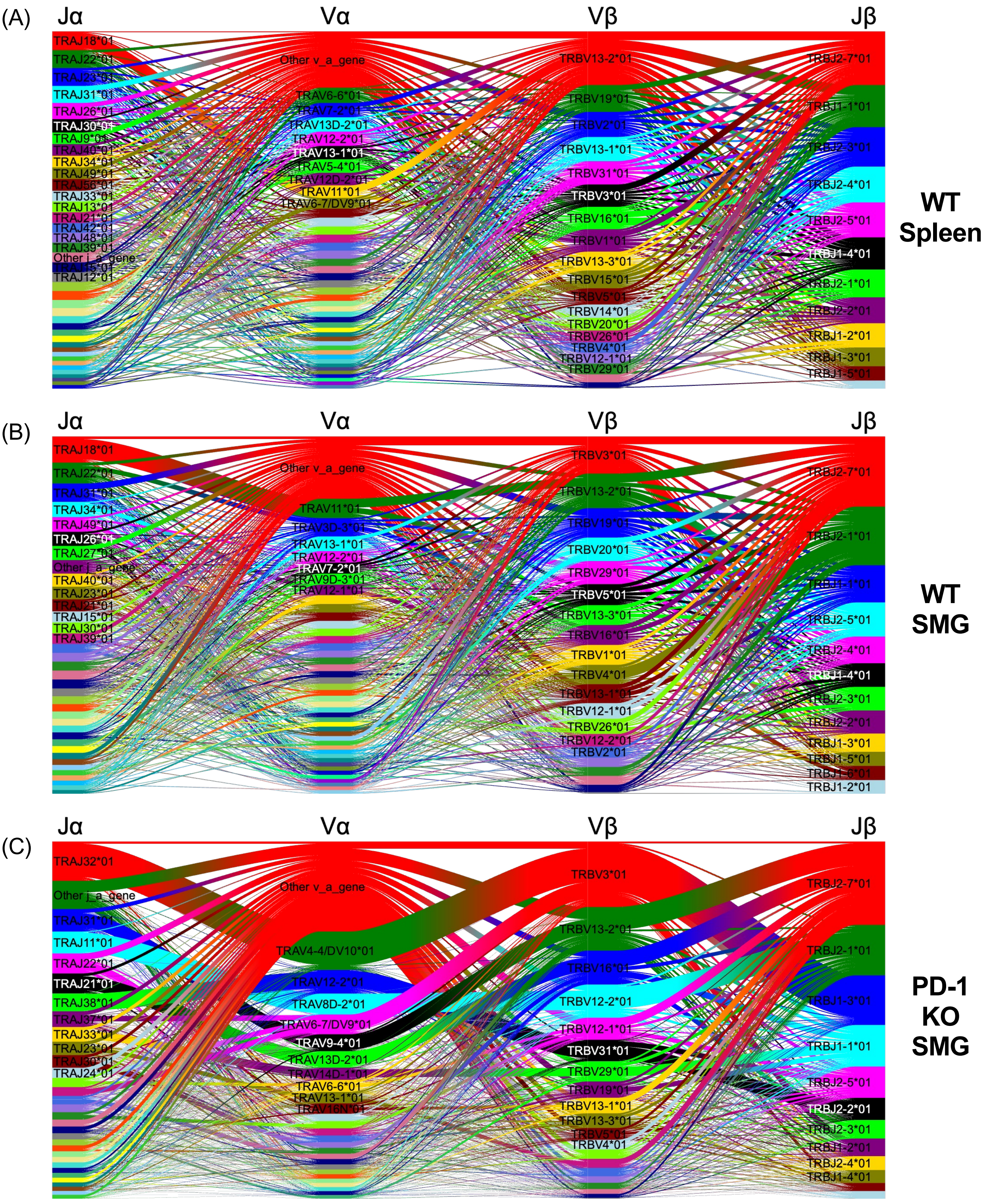


Supplemental Figure 6

Supplemental Table 1: Barcodes for hashtag and CITE-seq antibody panel. Samples were differentiated by barcoded hashtags, and cell surface antigens were quantified via barcoded CITE-seq antibodies.
feature_refs_totalseq_2023.csv

Supplemental Table 2: Sample specific metadata. Detailed information on samples sequenced, including age, sex, and hashtag number.
IGT15_sample_specific_metadata.csv

| | gene.x | xu_avg_log2FC | xu_p_val_adj | gene.y | borys_avg_log2FC | borys_p_val_adj | cluster.y |
|---|---|---|---|---|---|---|---|
| 1 | CCL5 | 1.1592174 | 0.00e+00 | Ccl5 | 2.0494649 | 0.000000e+00 | CD8 Activated 3 |
| 2 | CD27 | 0.7845661 | 1.04e-200 | Cd27 | 0.3943025 | 1.510004e-18 | CD8 Activated 3 |
| 3 | CD3E | 0.6235327 | 1.29e-223 | Cd3e | 0.5876111 | 4.068686e-197 | CD8 Activated 3 |
| 4 | CD3G | 0.4473554 | 9.01e-123 | Cd3g | 0.7267219 | 3.644860e-245 | CD8 Activated 3 |
| 5 | CD8A | 1.3385241 | 0.00e+00 | Cd8a | 1.3537279 | 0.000000e+00 | CD8 Activated 3 |
| 6 | CST7 | 0.4348996 | 3.96e-131 | Cst7 | 0.7543950 | 1.407018e-100 | CD8 Activated 3 |
| 7 | DUSP2 | 0.4524392 | 1.92e-105 | Dusp2 | 0.4928707 | 5.157834e-33 | CD8 Activated 3 |
| 8 | EOMES | 0.7727167 | 3.79e-262 | Eomes | 0.7497742 | 2.259535e-81 | CD8 Activated 3 |
| 9 | GIMAP5 | 0.6331413 | 1.80e-134 | Gimap5 | 0.3339846 | 3.670803e-15 | CD8 Activated 3 |
| 10 | GIMAP7 | 0.5359733 | 1.60e-121 | Gimap7 | 0.7291419 | 4.947802e-96 | CD8 Activated 3 |
| 11 | GZMA | 0.5218877 | 1.90e-125 | Gzma | 1.0987982 | 1.176317e-16 | CD8 Activated 3 |
| 12 | GZMK | 2.5444274 | 0.00e+00 | Gzmk | 1.6354820 | 0.000000e+00 | CD8 Activated 3 |
| 13 | HCST | 0.3838129 | 2.29e-98 | Hcst | 0.2786760 | 1.686570e-23 | CD8 Activated 3 |
| 14 | LCK | 0.4802215 | 1.09e-106 | Lck | 0.2808766 | 3.618570e-36 | CD8 Activated 3 |
| 15 | SH2D1A | 0.6189430 | 4.26e-144 | Sh2d1a | 0.8139796 | 2.917399e-93 | CD8 Activated 3 |
| 16 | TIGIT | 0.5071068 | 1.42e-130 | Tigit | 1.3033664 | 9.012945e-287 | CD8 Activated 3 |
| 17 | TRAC | 0.5911532 | 3.08e-108 | Trac | 0.5199020 | 5.784565e-51 | CD8 Activated 3 |
| 18 | CTSW | 0.3377142 | 7.25e-130 | Ctsw | 0.5715557 | 3.006485e-02 | CD8 Activated 1 |
| 19 | FYN | 0.4333480 | 6.93e-87 | Fyn | 1.5957326 | 1.279816e-30 | CD8 Activated 1 |
| 20 | ITGA4 | 0.4932247 | 1.94e-92 | Itga4 | 1.6522861 | 2.153123e-17 | CD8 Activated 1 |
| 21 | PTPRC | 0.3950021 | 1.21e-115 | Ptprc | 1.7468764 | 3.780634e-67 | CD8 Activated 1 |
| 22 | THEMIS | 0.4856210 | 8.67e-108 | Themis | 2.1907340 | 5.803560e-32 | CD8 Activated 1 |
| 23 | CNN2 | 0.5621824 | 4.84e-120 | Cnn2 | 0.3031852 | 3.051090e-12 | CD4 Activated 2 |
| 24 | CXCR3 | 0.8329156 | 0.00e+00 | Cxcr3 | 0.7415781 | 2.954837e-49 | CD4 Activated 2 |
| 25 | IL7R | 0.6186780 | 1.76e-119 | Il7r | 0.7606107 | 3.399001e-63 | CD4 Activated 2 |
| 26 | SIT1 | 0.5257278 | 1.44e-110 | Sit1 | 0.6527035 | 1.969976e-37 | CD4 Activated 2 |
| 27 | TCF7 | 0.5033040 | 4.54e-142 | Tcf7 | 0.7198954 | 6.851683e-55 | CD4 Activated 2 |
| 28 | EEF1A1 | 0.2656901 | 1.61e-126 | Eef1a1 | 0.2749972 | 5.127282e-32 | CD8 Activated 2 |
| 29 | KLRK1 | 0.6374698 | 3.90e-207 | Klrk1 | 0.6846949 | 2.447265e-48 | CD8 Activated 2 |
| 30 | RPS12 | 0.2947044 | 8.08e-133 | Rps12 | 0.3409599 | 1.290611e-15 | CD8 Activated 2 |
| 31 | RPS15A | 0.2921964 | 3.60e-142 | Rps15a | 0.3349600 | 2.085221e-51 | CD8 Activated 2 |
| 32 | RPS18 | 0.2867131 | 8.61e-120 | Rps18 | 0.5107942 | 2.948850e-83 | CD8 Activated 2 |
| 33 | RPS3 | 0.3127151 | 1.95e-139 | Rps3 | 0.3127010 | 6.446035e-47 | CD8 Activated 2 |
| 34 | LYAR | 0.9241965 | 0.00e+00 | Lyar | 0.2517994 | 9.202742e-13 | CD8 Proliferating |
| 35 | B2M | 0.2797617 | 9.36e-114 | B2m | 0.3181491 | 2.044477e-07 | CD8 IFN-Stim |
| 36 | CD84 | 0.4966092 | 1.07e-128 | Cd84 | 0.3838788 | 4.677757e-06 | Vγ6 Vδ4 |
| 37 | GIMAP4 | 0.5433050 | 2.44e-94 | Gimap4 | 0.5091128 | 1.897995e-25 | NKT |
| 38 | CD2 | 0.4521022 | 1.98e-100 | Cd2 | 0.9951835 | 9.383111e-05 | CD4 Tregs |
| 39 | KLRG1 | 0.4771995 | 4.39e-94 | Klrg1 | 1.1981061 | 1.118978e-13 | CD4 Tregs |

Supplemental Table 3: Overlapping differentially expressed genes between murine SMG T cell clusters and the human Sjögren's GzmK$^{+}$ CD8$^{+}$ T cell cluster. Differentially expressed genes specific to each cluster compared were identified with $log_2FC > 0.25$. The intersecting genes from each cluster compared to the top 50 differentially expressed genes specific to the GzmK$^{+}$ CD8$^{+}$ $T_M$ cluster from human Sjögren's Syndrome patients were identified.

### Graphical Abstract:

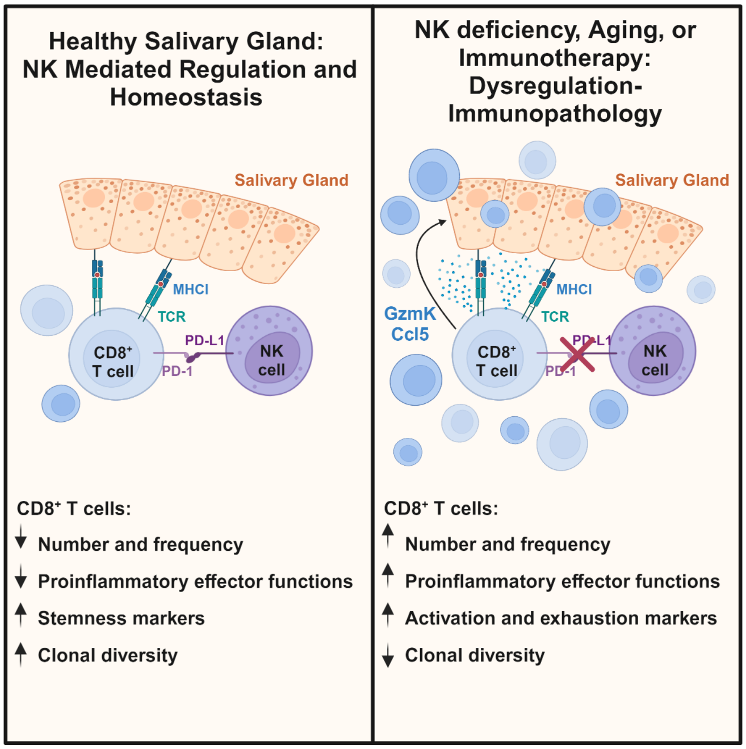
